# Ticks (Acari: Ixodida) on synanthropic small and medium-sized mammals in areas of the northeastern United States infested with the Asian longhorned tick, *Haemaphysalis longicornis*

**DOI:** 10.1101/2023.04.07.536052

**Authors:** Francisco C. Ferreira, Julia González, Matthew T. Milholland, Grayson A. Tung, Dina M. Fonseca

**Author notes:** These authors contributed equally to this work (Francisco C. Ferreira, Julia González).

## Abstract

The northeastern United States is a hot spot for tick-borne diseases. Adding to an already complex vector landscape, in 2017 large populations of the invasive *Haemaphysalis longicornis*, the Asian longhorned tick, were detected in New Jersey (NJ) and later found to be widespread from Connecticut to Georgia. In its native range in northeastern Asia, *H. longicornis* is considered an important vector of deadly pathogens to humans, companion animals, and livestock. To identify the primary hosts of *H. longicornis* we surveyed synanthropic small and medium-sized mammals in three different sites in suburban New Brunswick, NJ. Specifically, we collected approximately 9,000 tick specimens belonging to nine species from 11 different species of mammals sampled between May and September 2021. We found that *H. longicornis* feeds more frequently on rodents than previously thought, and that this invasive tick is likely exposed to important enzootic and zoonotic pathogens. Overall, we obtained detailed information about the seasonal dynamics and feeding patterns of six tick species common in the northeastern US, *Haemaphysalis longicornis*, *Amblyomma americanum, Dermacentor variabilis, Ixodes scapularis, I. texanus* and *I. cookei*. We found that unlike *I. scapularis* that feeds on mammals of all sizes, *H. longicornis* feeds on hosts following the general pattern of *A. americanum*, favoring larger species such as skunks, groundhogs, and raccoons. However, our survey revealed that unlike *A. americanum*, *H. longicornis* reaches high densities on Virginia opossum. Overall, the newly invasive *H. longicornis* was the most abundant tick species both on multiple host species and in the environment, raising significant questions regarding its role in the epidemiology of tick-borne pathogens, especially those affecting livestock, companion animals and wildlife. In conclusion, our findings provide valuable insights into the tick species composition on mammal hosts in New Jersey and the ongoing national expansion of *H. longicornis*.

## 1. Introduction

Tick-borne diseases affecting humans in the northeastern United States have been on the rise since 2000 (Rosenberg et al., 2018) primarily as a result of the geographical expansion of *Ixodes scapularis*, the blacklegged tick (Eisen et al., 2016), and the recolonization of previously occupied areas by *Amblyomma americanum*, the lone star tick (Rochlin et al., 2022). These tick species are efficient vectors of multiple viral, bacterial and protozoan pathogens, many of which were described in the last 20 years infecting humans, companion animals and wildlife (Goddard and Varela-Stokes, 2009; Tokarz et al., 2018; Madison-Antenucci et al., 2020; Fleshman et al., 2022).

While some tick species are considered critical pathogen vectors, their importance varies seasonally and with the relative abundance of different reservoir they use as hosts, which in turn determines the dynamics of animal-to-animal (enzootic) and animal-to-human (zoonotic) transmission cycles (Krasnov et al., 2007; Linske et al., 2018; Ginsberg et al., 2021). Thus, it is essential to evaluate associations between tick species and their vertebrate hosts to assess pathways of tick-borne pathogen transmission. This is especially the case when trying to evaluate the potential of an invasive tick species as a vector of pathogens in its new range.

The invasive Asian longhorned tick, *Haemaphysalis longicornis*, was detected for the first time in the western hemisphere in 2017 in New Jersey, northeastern USA (Rainey et al., 2018), and it has since been detected in 18 states (USDA, 2023). This species reaches particularly high densities in transitional habitats between meadows or pathways and forests, which are areas intensely used by humans and dogs (González et al., 2023). In its native range in northeastern China, Korea, and Japan, *H. longicornis* is considered a critical pathogen vector of *Dabie bandavirus*, the causative agent of human severe fever with thrombocytopenia syndrome (Casel et al., 2021). In Asia, it is also a significant vector of agents causing canine and livestock babesiosis (Shaw et al., 2001; Bai et al., 2002; Guan et al., 2010; Sivakumar et al., 2014), and in Australasia, where it became established in the early 20^th^ century, it vectors *Theileria orientalis,* the agent of deadly theileriosis to cattle, resulting in very significant economic losses (Watts et al., 2016). This widespread distribution and adaptability of *H. longicornis* underscores the necessity of understanding its phenology and host associations, which could reveal potential impacts on human and animal health in new areas it colonizes.

Although the public health impact of this invasive tick species is still being investigated in the US, American *H. longicornis* are competent vectors of *Rickettsia rickettsii*, the bacterial agent of Rocky Mountain spotted fever (Stanley et al., 2020), and Heartland virus (Raney et al., 2022). In the US state of Virginia, deaths of cattle due to infection with *Theileria orientalis* Ikeda, a strain new to the US, have been attributed to transmission by *H. longicornis* (Dinkel et al., 2021). Additionally, a recent examination of field collected *H. longicornis* found larvae, nymphs and adults infected with Bourbon virus (Cumbie et al., 2022). Associations with other pathogens as populations expand across favorable areas are expected (Rochlin et al., 2023).

Within the northeastern US, New Jersey, the US state with the highest human density and urbanization (Bureau, 2021), stands out for its high frequency of human-tick encounters and high incidence of tick-borne diseases in humans and their companion animals (Bowman et al., 2009; Nieto et al., 2018; Mahachi et al., 2020). Our objective was to identify important mammalian hosts of *H. longicornis*. To do so we removed ticks from a variety of small and medium-sized mammals captured near paths and edges of forests where questing *H. longicornis* are most abundant (González et al., 2023). In the process we also collected the other epidemiologically important tick species in the NE US, allowing us to develop side-by-side comparisons of tick-host associations. Here we summarize our findings and discuss their implications to the epidemiology of tick-borne diseases in New Jersey and elsewhere.

## 2. Materials and methods

### 2.1. Study areas

We trapped small and medium-sized mammals and conducted environmental tick sampling in three sites within the Rutgers University Cook Campus in New Brunswick, New Jersey (Fig. 1), between May and September 2021. The “Goat Farm” (40.47302, −74.43727) is an area of approximately 0.13 km^2^ composed of small forest fragments and grass patches used for goat and sheep ranching, and is part of the Rutgers NJ Agricultural Experiment Station (NJAES); the “Rutgers Gardens” (40.47417, −74.42264) is a 0.73 km^2^ botanic garden composed of small forest patches and vegetable farms that receives around 35,000 visitors per year; the “University Inn” (40.48394, - 74.43020) is a 0.06 km^2^ meadow and forested park behind the Rutgers University Inn & Conference Center. Oak trees and huckleberry shrubs dominate forest composition in all sampling areas. We chose these areas because they provided a variety of different habitats, which we hoped would maximize the diversity and number of mammals we might be able to survey. In addition, previous work indicated the presence from spring to fall of significant populations of *H. longicornis*, *I. scapularis* and *A. americanum* (González et al. 2023), our focal tick species and the two most significant vectors of tick-borne disease agents in NJ (Madison-Antenucci et al., 2020).

**Fig. 1.**
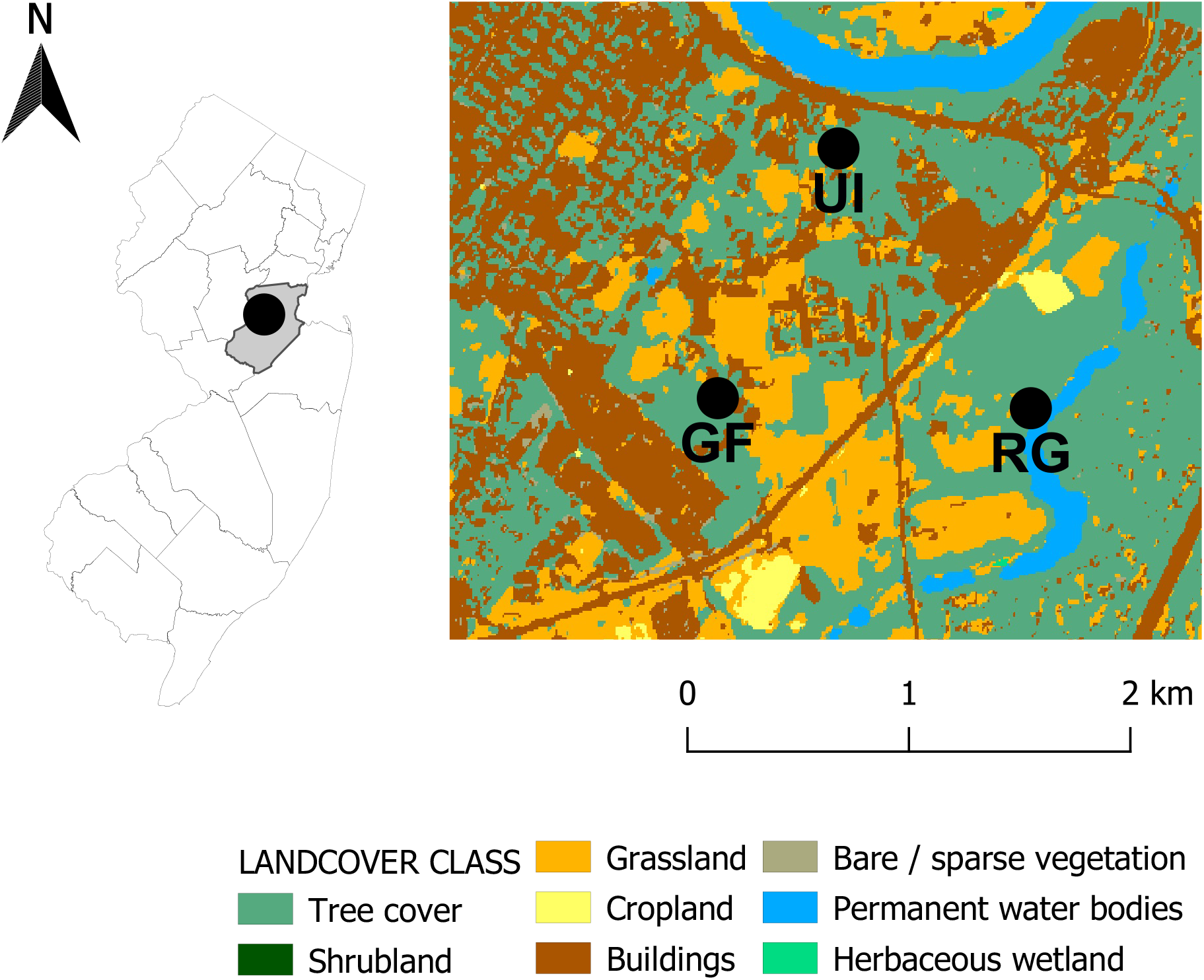
Map of the trapping sites within the Rutgers University Cook Campus in New Brunswick NJ, USA, 2021. Goat Farm (GF), Rutgers Gardens (RG) and University Inn (UI). QGIS map base ESA World Cover (Zanaga et al., 2021).

### 2.2. Mammal trapping and processing

We set up traps at dusk each day and checked them for the first time the following dawn. To capture day-active mammals, we kept all traps open, checking them at least hourly until noon or until temperatures neared 27°C (we are referring to this combination of night and daylight sampling as a “1 trap night-day”). Each sampling event lasted 3 consecutive trap night-days, and we sampled each site 4 times from May 12 to September 15. In total, we trapped mammals at each site for 12 trap night-days.

Depending on the area and the suitability of the vegetation of each site, we set up 30 ± 7 (Mean ± SD) Sherman traps (8 × 9 × 23 cm, H.B. Sherman Traps, Tallahassee, FL, USA), 14 ± 4.2 medium Tomahawk traps (13 × 13 × 40 cm, Tomahawk Live Trap, Hazelhurst, WI, USA) and 11 ± 1.4 large Tomahawk traps (30 × 30 × 81 cm, Tomahawk Live Trap, Hazelhurst, WI, USA) each night. We baited Sherman and medium Tomahawk traps with fresh apple slices and a mix of peanut butter, oats, and vanilla extract. We provided Sherman traps with cotton balls to improve mammal well-being. We baited large Tomahawk traps with either fresh produce (apple, carrots, and cantaloupe) or with sardines and wet cat food. We placed Sherman traps at ∼10 m intervals along areas of transition between meadows and forests (i.e., ecotones; see González et al. 2023) and areas of interface between shrubby or tall grass areas and open meadows. We also placed Sherman traps inside forest fragments during the first and second trapping weeks on each site. However, due to zero captures we did not do so during the third and fourth trapping events. We placed medium and large Tomahawk traps interspersed with the Sherman traps within forest fragments and near trashcans and dumpsters, which are often hot spots of activity for skunks and other medium-sized mammals (Bozek et al., 2007).

We anesthetized mammals trapped in Sherman traps (house mice, *Mus musculus*; *Peromyscus* mice; meadow voles, *Microtus pennsylvanicus*; short-tailed shrews, *Blarina brevicauda* and eastern chipmunks, *Tamias striatus*) and medium Tomahawk traps (eastern chipmunks and eastern gray squirrels, *Sciurus carolinensis,* henceforth gray squirrels) with isoflurane (99.9%, Decra, Overland Park, KS, USA). We anesthetized larger animals such as eastern cottontail rabbit (*Sylvilagus floridanus*), raccoons (*Procyon lotor*), Virginia opossums (*Didelphis virginiana*) striped skunks (*Mephitis mephitis*), and groundhogs (*Marmota monax*) with intramuscular injection of ketamine hydrochloride (100 mg/mL, Decra, Overland Park, KS, USA) and dexmedetomidine (0.5 mg/mL, Decra, Overland Park, KS, USA) mixed in the syringes, adjusting the final doses depending on the weight and animal species. We used metal rods and trap dividers (Heavy duty Divider for 12" traps, Tomahawk Live Trap, Hazelhurst, WI, USA) to restrain the animals to one side of the cage to inject them with the anesthetics. When deep anesthesia levels were not reached, we administered an additional half dose of the anesthetic mix. We used Atipemazole (5mg/mL, Modern Veterinary Therapeutics, Sunrise, FL, USA) injected intramuscularly to a dose five times higher than the administered dexmedetomidine dose to reverse the anesthesia. This anesthesia protocol allowed us to process the animals safely for 30-50 min. We released all animals in the same site they were captured after they recovered full alertness and movement capacities.

We inspected smaller mammals (< 150 g average weight; all animals captured in Sherman traps) carefully for at least 5 min, which is the minimum recommended time period (Estrada-Peña et al., 2021), to reduce the chances of false-negative results in tick detection. We examined medium-sized mammals for ticks for at least 10 min, although in most cases the inspection took between 30-50 min. We collected and kept only ticks that were attached to the hosts and stored them in tubes containing 100% ethanol with the aid of thin forceps. When tick abundance was too high, we collected representative subsamples of ticks from all body parts. We did not process animals recaptured within the same sampling event (i.e., 3 consecutive trap night-days), instead we noted the recapture and released them. These procedures were approved by Rutgers University Institutional Animal Care and Use Committee (PROTO201900163) and were conducted under NJ Division of Fish and Wildlife permit number SC 2021-01.

### 2.3. Questing tick sampling

We conducted environmental tick sampling between 09:00 h and 16:00h at each site one week before or one week after mammal trapping campaigns at the same site, and once each in October and November of 2021. We completed a total of 11 sessions of tick sampling per site, flagging between 50-75 m^2^ per session in different habitats where we had or would set up the mammal traps. We collected ticks from the ground cover with modified tick sweeps (Egizi et al., 2019a) that consisted of a white crib flannel (0.5 x 1 m, Shunjie.Home, Shandong, China) attached to the short arm of a L-shaped PVC pipe. We checked the sweep cloth for the presence of ticks every 1-2 m to avoid tick drop-off over longer intervals during the sampling (Bickerton et al., 2021). We stored nymphs and adult ticks in screwcap 50 ml tubes and removed larvae from the sweeps with transparent adhesive tape. We counted and identified all ticks in the laboratory, where they were kept at −80°C before and after identification.

### 2.4. Tick identification

We identified tick species using taxonomic keys that include species from the Northeastern regions of the United States (Durden and Keirans, 1996; Keirans and Durden, 1998; Egizi et al., 2019b). On some occasions, we confirmed tick identification via molecular barcoding, which entails amplifying and sequencing a ∼ 400-700 bp fragment of the mitochondrial cytochrome oxidase I gene (*cox*1). For that, we extracted DNA from nymph or adult legs or from entire tick larvae using a “HotSHoT” (hot sodium hydroxide and Tris) method (Johnson et al., 2015). If the specimen was a putative *H. longicornis,* we used primer pairs cox1F and cox1R (Chitimia et al., 2010; Egizi et al., 2020). For other tick species, especially those belonging to the genus *Ixodes*, we used primers LCO1490 and HCO2198 (Folmer et al., 1994). For *Ixodes marxi,* because the standard primers used for DNA barcoding were not working, we designed and used a new set of primers (Imarxi16F – 5’ TTTTGGGAGATGATCGGCC 3’ and Imarxi579R – 5’ AGTATGGTAATTGCTCCTGC 3’) targeting the *cox*1 gene of the voucher *I. marxi* sequenced and deposited by Ondrejicka et al. (2017). PCR products were visualized in a 1% agarose gel, positive samples were purified with ExoSAP-IT (USB Corporation, Cleveland, OH, USA) following the manufacturer’s protocol, and the products were directionally sequenced by Genewiz (Piscataway, NJ, USA). We quality-trimmed sequences with Geneious Prime 2023.0.4 (Biomatters Inc. San Diego, CA, USA) and the consensus was used as a query in NCBI’s Basic Local Alignment Search Tool, BLASTn (Altschul et al., 1990). Nucleotide sequence data related to the molecular identification of ticks are available in the GenBank database under the accession numbers: OQ690118-OQ690122.

### 2.5. Statistical analyses

We calculated the Shannon diversity index (Shannon, 1948), considering the number of specimens of each species in all sites using the *vegan* package (Oksanen et al., 2020) in the statistical software R (R Development Core Team, 2020). Then, we compared the animal diversity among sites using the Hutcheson t-test (Hutcheson, 1970), a formula developed to compare the diversity of two community samples using the Shannon index (Zar, 2010). In addition, we evaluated the overall trapping success among sites, and among different types of traps (Sherman, medium and large Tomahawk) within the same site using Yates’s chi-square tests.

We assessed the association among ticks and mammal species by computing the infestation prevalence of each tick species and the abundance of each tick species and stage on each mammal species. We calculated prevalence by dividing the number of tick infested hosts by the number of hosts processed, and calculated tick intensity and tick abundance by dividing the total number of ticks by the number of infested hosts and by the total number of mammals of each species processed, respectively (Margolis et al., 1982; Bush et al., 1997). In addition, we used the hybrid Wilson/Brown method to calculate 95% confidence intervals (CI) of the proportion of animal infested by each tick species to assess the precision of these results (Brown et al., 2001).

We checked for differences in tick abundance among male and female hosts of each mammal species using Mann-Whitney tests, except when distributions were normally distributed, such as for meadow voles and striped skunks, to which we applied a t-test with Welch’s correction. We used non-parametric Kruskal-Wallis test and Dunn’s multiple comparisons tests to evaluate differences in ranges of abundance of the most prevalent tick species in mammalian hosts (those that infested over 10% of the animals captured) per life stage. We used Spearman correlations to assess whether mean weight of mammals captured was associated with tick abundance by analyzing each life stage separately. We performed simple linear regressions when significant correlations were detected. Because there was significant variation in the species and numbers of mammals caught in each of the three sites across trapping events we combined the data on tick occurrence from all sites.

To normalize the distribution of the number of questing ticks collected from the environment (ticks/m^2^) we (Ln (Y+1)) transformed the data. We examined differences in relative abundance between sites and months using two-way ANOVA and Tukey honestly significant difference (HSD) tests. In addition, we compared the relative abundance of each tick species among sites using ANOVA tests. All statistical analysis and graphs were executed using the software GraphPad Prism version 8 GraphPad Software, San Diego, California USA).

## 3. Results

### 3.1 Trapping effort and mammal composition comparisons among sites

We captured 329 mammals belonging to 11 species with an effort of 2,194 trap night-days and an overall trapping success rate of 15% (Table 1). We processed 273 mammals because 51 individuals were recaptured during the same sampling event and therefore not re-processed and five escaped before being processed. Twenty-four of these 273 mammals were captured 1-3 times again during other sampling events with time between captures ranging from 20 to 104 days (mean = 44 days). This intervening time was enough to erase the effects of our previous tick removal because Ixodidae ticks commonly feed to repletion on mammals within 20 days (Sonenshine and Roe, 2013). Therefore, these repeated captures were processed and analyzed as separate events. Most mammals were adults (260/273), 48% males and 47% females. A few juveniles (n = 13) could not be sexed.

**Table 1.** Assemblage descriptors of small and medium-sized mammals captured at three sites in New Brunswick NJ, USA, 2021: Number of individuals captured (n), number of trap/nights (TN), trapping success (%), number of processed animals and species richness (S). We calculated the Shannon index to estimate the animal diversity (H’) and compared it between sites using Hutcheson t-tests. Differences in trapping success among sites were calculated using Yates’s chi-square tests. Different letters show significant differences (P < 0.05).

| Site | Sherman |  |  | Medium Tomahawk |  |  | Large Tomahawk |  |  | Total traps |  |  | Processed animals | Species Richness (S) | Shannon Index (H') |
| --- | --- | --- | --- | --- | --- | --- | --- | --- | --- | --- | --- | --- | --- | --- | --- |
|  | n | TN | Trapping Success | n | TN | Trapping Success | n | TN | Trapping Success | n | TN | Trapping Success |  |  |  |
| Goat Farm | 123 | 424 | 29.0<sup>a</sup> | 27 | 191 | 14.1<sup>a</sup> | 37 | 150 | 24.7<sup>a</sup> | 187 | 765 | 24.4<sup>a</sup> | 147 | 11 | 1.817<sup>a</sup> |
| Rutgers Gardens | 46 | 353 | 13.0<sup>b</sup> | 23 | 174 | 13.2<sup>a</sup> | 18 | 134 | 13.4<sup>b</sup> | 87 | 661 | 13.2<sup>b</sup> | 80 | 8 | 1.839<sup>a</sup> |
| University Inn | 9 | 504 | 1.8<sup>c</sup> | 19 | 138 | 13.8<sup>a</sup> | 27 | 126 | 21.4<sup>ab</sup> | 55 | 768 | 7.2<sup>c</sup> | 46 | 8 | 1.641<sup>a</sup> |
| **Total** | **178** | **1281** | **13.9** | **69** | **503** | **13.7** | **82** | **410** | **20.0** | **329** | **2194** | **15.0** | **273** | **11** | **2.025** |

Hutcheson t-tests (P > 0.05): Goat Farm vs Rutgers Gardens $t_{262} = 0.22$ ; Goat Farm vs University Inn $t_{121} = 1.36$ ; Rutgers Gardens vs University Inn $t_{97} = 1.62$ . Yates's chi-square tests for total traps: Goat Farm vs Rutgers Gardens $\chi^2 = 28.36$ (P < 0.0001); Goat Farm vs University Inn $\chi^2 = 84.82$ (P < 0.0001); Rutgers Gardens vs University Inn $\chi^2 = 13.63$ (P < 0.001). Large Tomahawk: Goat Farm vs Rutgers Gardens $\chi^2 = 5.02$ (P < 0.05); Goat Farm vs University Inn $\chi^2 = 0.24$ (P > 0.05); Rutgers Gardens vs University Inn $\chi^2 = 2.37$ (P > 0.05). Medium Tomahawk (all P > 0.05): Goat Farm vs Rutgers Gardens $\chi^2 = 0.01$ ; Goat Farm vs University Inn $\chi^2 = 0.004$ ; Rutgers Gardens vs University Inn $\chi^2 = 0.0007$ . Sherman (all P < 0.0001): Goat Farm vs Rutgers Gardens $\chi^2 = 27.96$ ; Goat Farm vs University Inn $\chi^2 = 137.7$ ; Rutgers Gardens vs University Inn $\chi^2 = 41.86$ .

The highest trapping success rate was at Goat Farm, followed by Rutgers Gardens and University Inn (Table 1). We found the greater species richness at Goat farm, and that was the site where we also captured the highest number of mammals, with *Peromyscus* mice as the most common species (Fig. 2). In Rutgers Gardens, we captured a high number of eastern chipmunks (Fig. 2). We captured the overall lowest number of animals from the University Inn site (Table 1), with a trapping success of the Sherman traps significantly lower than that of the medium (X2 = 34.47, P < 0.0001) and large Tomahawk traps (X2 = 68.59, P < 0.0001). Although the overall mammal diversity was not different among sites based on the Shannon’s diversity index (Table 1), there was an absence of house mice and short-tailed shrews and almost no *Peromyscus* mice and meadow voles at the University Inn (Fig. 2). The very low numbers of small mammals are reflected in the lower success rate for the Sherman traps at the University Inn compared to the other sites (Table 1) and indicate potentially significant differences in the mammal community composition.

**Fig. 2.**
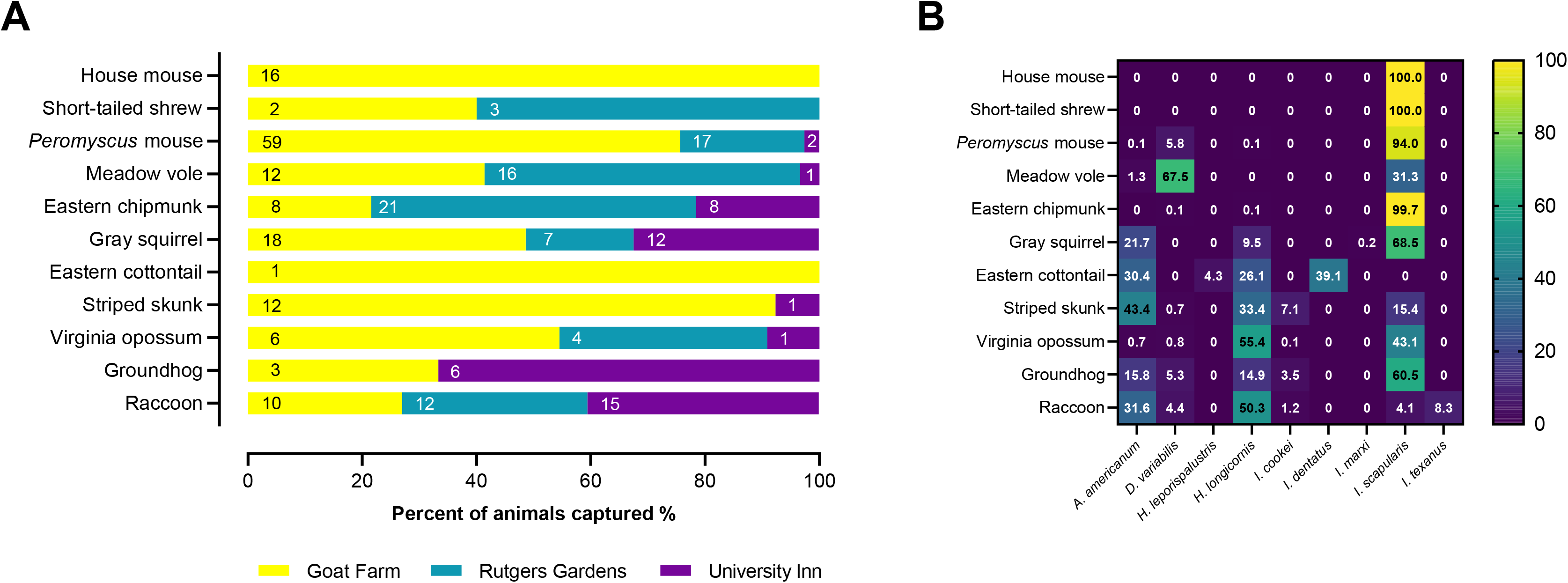
A) Animal diversity found at each trapping site in New Brunswick NJ, USA, 2021. Numbers inside bars represent the number of individuals captured at each site. B) Heat map of tick diversity collected from each host species captured. Values in the cells are the percentages that each tick species represents among all ticks collected from each host species.

### 3.2. Ticks on hosts

We collected a total of 9,111 ticks from the mammals processed, and 9,009 were identified into nine different species (Table 2). Overall, 89% (243/273) of the animals were infested with at least one tick, including all raccoons, groundhogs, Virginia opossums and the eastern cottontail rabbit (Supplementary Table S1). *Ixodes scapularis* was the most prevalent species (77.6%), followed by *A. americanum* (31.1%), *H. longicornis* (27.8%) and *D. variabilis* (22.8%); the remaining tick species showed a prevalence lower than 10% (Supplementary Table S1). Four tick species were found on one mammal species only: *Ixodes texanus* on raccoons, and *Haemaphysalis leporipalustris* and *Ixodes dentatus* on the single eastern cottontail rabbit we examined and *Ixodes marxi* (new species record for NJ) on two gray squirrels. The two 430 bp *cox1* sequences obtained, GenBank acc. num. OQ690118, were identical and had 100% identity to a female *I. marxi* collected in Canada (Ondrejicka et al., 2017). *Ixodes scapularis* was collected from 10 out of 11 different mammal species, while *A. americanum, D. variabilis, H. longicornis* and *I. cookei* infested from four to eight mammal species. We found that *A. americanum* and *H. longicornis* showed a similar host range, infesting mostly larger animals (Fig. 2).

**Table 2.** Total number of mammals examined and ticks collected (adults; nymphs; larvae) per host species in New Brunswick NJ, USA, 2021.

| Animal species | n | *Amblyomma americanum* | *Dermacentor variabilis* | *Haemaphysalis leporispalustris* | *Haemaphysalis longicornis* | *Ixodes cookei* | *Ixodes dentatus* | *Ixodes marxi* | *Ixodes scapularis* | *Ixodes texanus* | Total ticks |
| --- | --- | --- | --- | --- | --- | --- | --- | --- | --- | --- | --- |
| house mouse | 16 | 0; 0; 0 | 0; 0; 0 | 0; 0; 0 | 0; 0; 0 | 0; 0; 0 | 0; 0; 0 | 0; 0; 0 | 0; 5; 29 | 0; 0; 0 | 34 |
| short-tailed shrew | 5 | 0; 0; 0 | 0; 0; 0 | 0; 0; 0 | 0; 0; 0 | 0; 0; 0 | 0; 0; 0 | 0; 0; 0 | 0; 1; 40 | 0; 0; 0 | 41 |
| *Peromyscus* mouse | 78 | 0; 0; 1 | 0; 12; 49 | 0; 0; 0 | 0; 0; 1 | 0; 0; 0 | 0; 0; 0 | 0; 0; 0 | 0; 106; 881 | 0; 0; 0 | 1050 |
| meadow vole | 29 | 0; 0; 1 | 0; 19; 35 | 0; 0; 0 | 0; 0; 0 | 0; 0; 0 | 0; 0; 0 | 0; 0; 0 | 0; 8; 17 | 0; 0; 0 | 80 |
| eastern chipmunk | 37 | 0; 0; 0 | 0; 0; 1 | 0; 0; 0 | 0; 0; 1 | 0; 0; 0 | 0; 0; 0 | 0; 0; 0 | 0; 303; 370 | 0; 0; 0 | 675 |
| gray squirrel | 37 | 0; 68; 32 | 0; 0; 0 | 0; 0; 0 | 0; 2; 42 | 0; 0; 0 | 0; 0; 0 | 1; 1; 0 | 0; 91; 225 | 0; 0; 0 | 462 |
| eastern cottontail | 1 | 0; 4; 3 | 0; 0; 0 | 0; 1; 0 | 0; 0; 6 | 0; 0; 0 | 0; 9; 0 | 0; 0; 0 | 0; 0; 0 | 0; 0; 0 | 23 |
| striped skunk | 13 | 0; 96; 485 | 8; 2; 0 | 0; 0; 0 | 3; 100; 344 | 4; 27; 64 | 0; 0; 0 | 0; 0; 0 | 0; 19; 187 | 0; 0; 0 | 1339 |
| Virginia opossum | 11 | 0; 1; 8 | 10; 0; 0 | 0; 0; 0 | 25; 67; 633 | 0; 0; 1 | 0; 0; 0 | 0; 0; 0 | 0; 43; 521 | 0; 0; 0 | 1309 |
| groundhog | 9 | 0; 14; 4 | 6; 0; 0 | 0; 0; 0 | 0; 4; 13 | 1; 3; 0 | 0; 0; 0 | 0; 0; 0 | 0; 11; 58 | 0; 0; 0 | 114 |
| raccoon | 37 | 1; 300; 926 | 171; 0; 0 | 0; 0; 0 | 70; 232; 1652 | 1; 38; 9 | 0; 0; 0 | 0; 0; 0 | 1; 18; 141 | 75; 119; 128 | 3882 |
| **total** | **273** | **1; 483; 1460** | **195; 33; 85** | **0; 1; 0** | **98; 405; 2692** | **6; 68; 74** | **0; 9; 0** | **1; 1; 0** | **1; 605; 2469** | **75; 119; 128** | **9009** |

We collected one *H. longicornis* larva feeding on an eastern chipmunk and another one on a *Peromyscus* mouse. We checked identification by sequencing the *cox*1 barcoding locus and found in both cases 100% identity to the parthenogenetic *H. longicornis* H1 haplotype (GenBank acc. num. OQ690119-OQ690120), which is the most common in New Brunswick, NJ (Egizi et al., 2020). *Haemaphysalis longicornis* was the predominant species on Virginia opossums and raccoons (over 50% of total ticks collected from these hosts), while *I. scapularis* was predominant in six mammal species (Figs. 2 and 3). Indeed, more than 90% of the ticks collected from *Peromyscus* and house mice, short-tailed shrews, and eastern chipmunks were *I. scapularis*. However, immature *D. variabilis* were dominant on meadow voles (Figs. 2 and 3). Striped skunks and raccoons were highly parasitized by *A. americanum*, and these two mammals in addition to Virginia opossum were also the main hosts for *H. longicornis* (Fig. 4). The number of immature stages of *H. longicornis* and *A. americanum* collected from these hosts was significantly higher compared to other hosts (Supplementary Table S2). Raccoons were the hosts with the highest abundance of adult *D. variabilis*, although immature stages were collected mostly from meadow voles and *Peromyscus* mice (Table 2). The highest intensity and abundance of *I. scapularis* was found on Virginia opossums, although this tick species was common on almost all mammals (Fig. 3, Fig. 4 and Supplementary Table S1).

**Fig. 3.**
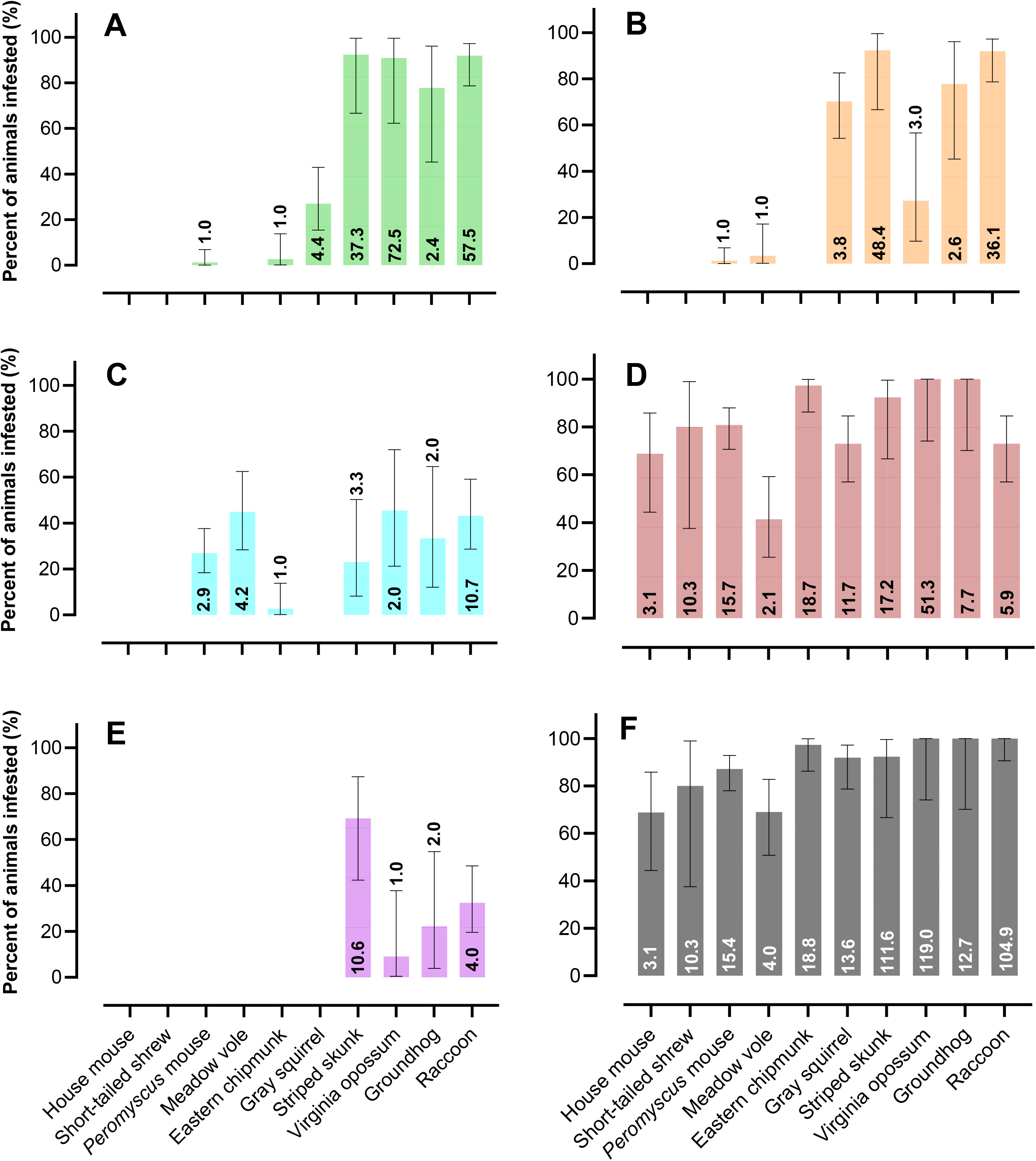
Infestation prevalence expressed as percentage (with 95% CI) of ticks from each host species in New Brunswick NJ, USA, 2021. A) *Haemaphysalis longicornis,* B) *Amblyomma americanum*, C) *Dermacentor variabilis*, D) *Ixodes scapularis*, E) *Ixodes cookei*, F) total number of identified ticks combined. Numbers inside or above bars represent tick intensity. Eastern cottontail is not included because only one individual was captured (see Supplementary Table S1 for more details).

**Fig. 4.**
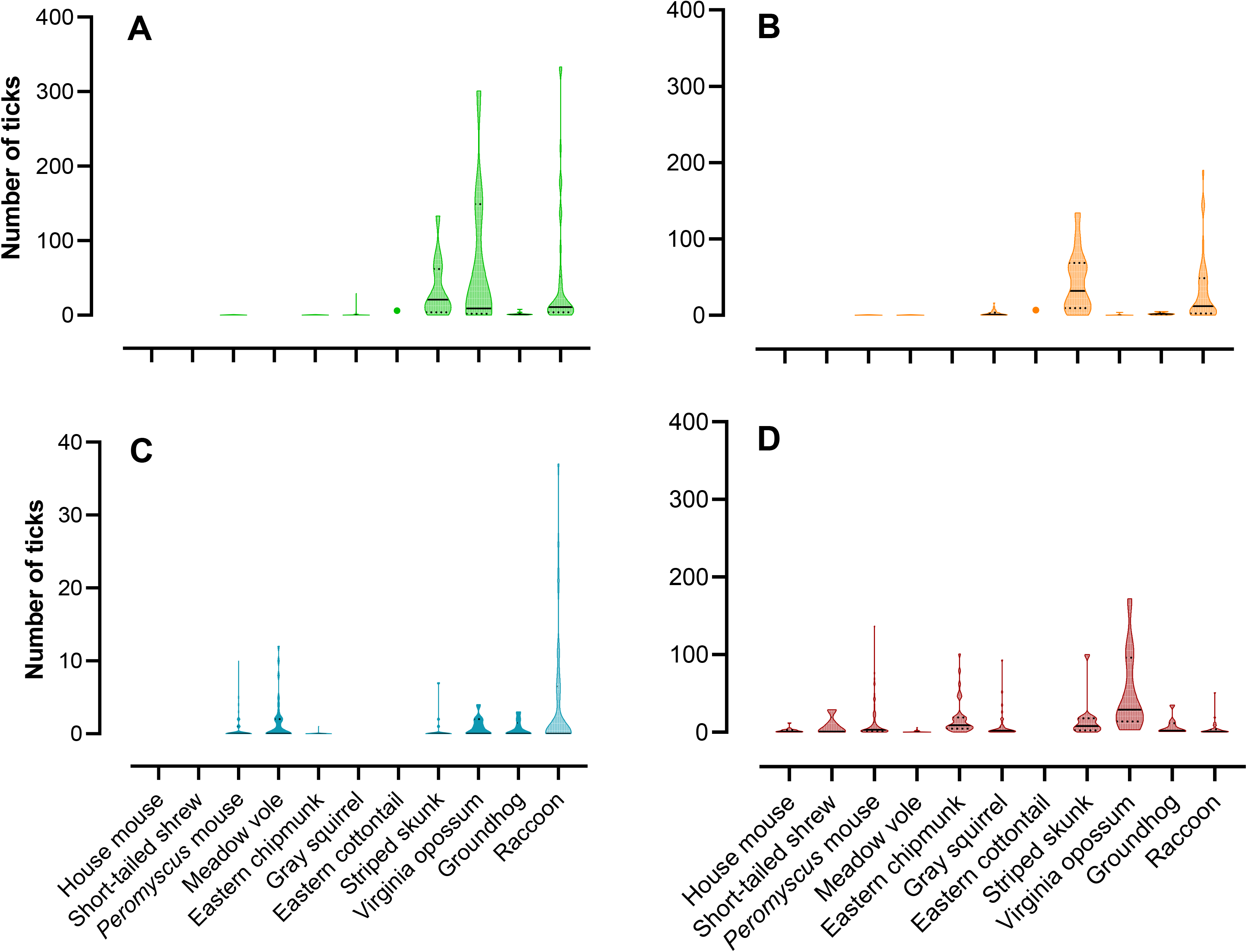
Number of individuals of each the four most prevalent tick species collected from all the host species trapped in New Brunswick NJ, USA, 2021. A) *Haemaphysalis longicornis*, B) *Amblyomma americanum*, C) *Dermacentor variabilis* (note the different Y axis scale), and D) *Ixodes scapularis*. In addition to showing the data distribution, the median is shown as a line and the quartiles as dotted lines.

Interestingly, the number of *I. scapularis* nymphs collected from eastern chipmunks was significantly higher than that found on other small mammals, such as *Peromyscus* and house mice, and meadow voles (Supplementary Table S2). Tick abundance varied with the size of the host species (Supplementary Table S3), but there was no sex bias (P > 0.05). The abundance of all stages of *H. longicornis*, larvae and nymphs of *A. americanum*, and adult *D. variabilis* was significantly (P < 0.001) positively correlated with the mean weight of the hosts (Supplementary Table S3). Overall, the larger animals (striped skunks, Virginia opossums, groundhogs, and raccoons) had the most ticks. However, this association was absent for all stages of *I. scapularis* (Supplementary Material Fig. S1).

The majority of *H. longicornis* adults were collected in late July and August, with most larvae found in August and September. *Haemaphysalis longicornis* nymphs were collected throughout the entire surveillance period although we found the highest numbers in June and July. We collected most *A. americanum* nymphs in May and June, while larvae were abundant from July to September. *Ixodes scapularis* nymphs were collected throughout the entire surveillance, with the highest numbers in May and June, while larvae were collected mostly in July and August (Fig. 5). Of note, we collected only a single adult *A. americanum* and one adult *I. scapularis* from the captured mammals (Table 2).

**Fig. 5.**
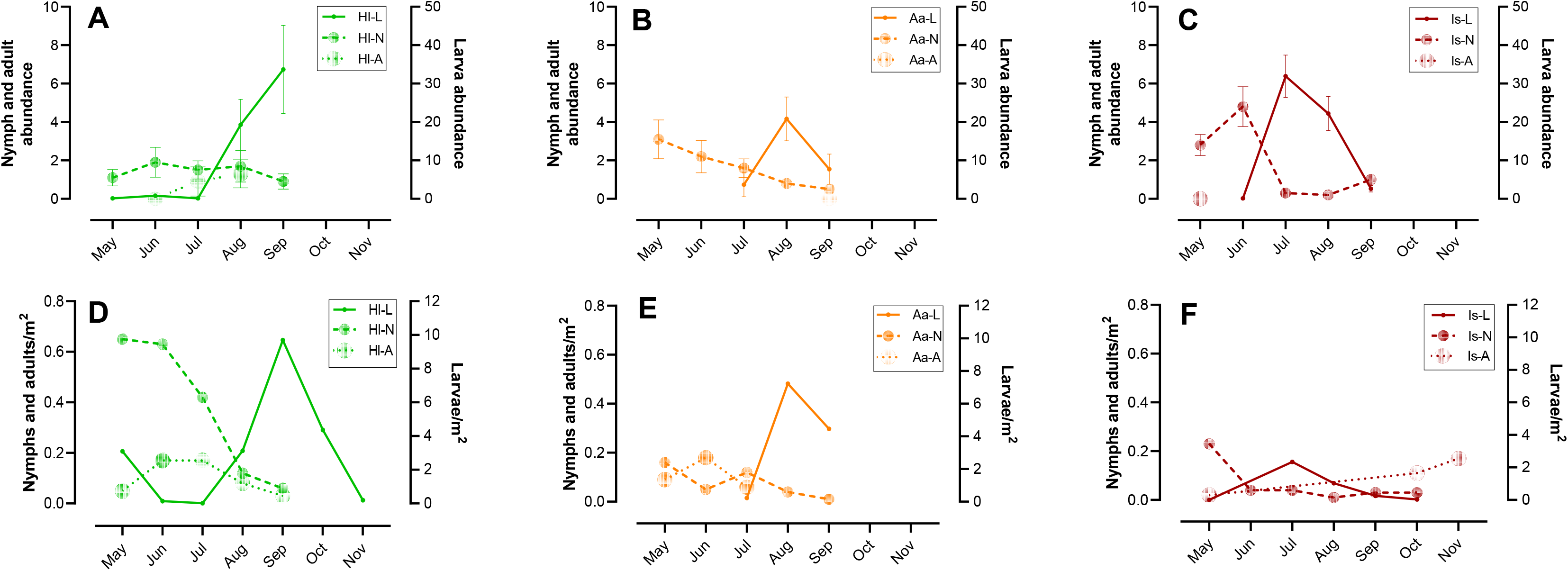
Seasonal dynamics of life stages of, *Haemaphysalis longicornis (Hl* = green), *Amblyomma americanum (Aa* = orange), and *Ixodes scapularis* (*Is*= red); L = larva, N = nymph, A = adult. A-C refer to ticks collected from animals captured and D-F refer to questing ticks by square meter sampled from the environment. If zero ticks were collected, they are not represented.

### 3.3. Questing ticks

In total, 10,836 ticks from five different species were collected by flagging, to an overall density of 5.6 ticks/m^2^ (Supplementary Table S4). Relative abundance of questing ticks varied according to the month of sampling, which explained 70% of the total variation (F_6,12_ = 7.39, P = 0.0017), but did not differ among sites (11% total variation; F_2,12_ = 3.48, P = 0.0642) (Supplementary Table S5). *Haemaphysalis longicornis* was the most abundant tick (42.5%; 2.4 ticks/m^2^), followed by *A. americanum* (35.6%; 2.0 ticks/m^2^) and *I. scapularis* (21.9%; 1.2 ticks/m^2^), while *D. variabilis* and *H. leporispalustris* constituted less than 0.1% of the ticks collected from the environment. We sequenced one nymph and one larva to confirm the identity of two specimens of *H. leporispalustris* (GenBank acc. num. OQ690121-OQ690122), which had 99.7% and 98.6% identity, respectively, with specimens collected in New Jersey and Virginia (Thompson et al., 2020). The abundance of each of the three most common tick species was also similar among sites (P > 0.05; *A. americanum* F = 0.1665; *H. longicornis* F = 2.4170; *I. scapularis* F = 0.0923).

The seasonal activity of *H. longicornis* nymphs was higher in May and June, gradually decreasing throughout September, after which they were no longer detected. Adults were more active in June and July, before the peak activity of larvae observed in September. The activity of questing adult *A. americanum* was higher in June, between the two nymphal peaks observed in May and July, with larval activity in August and September. *Ixodes scapularis* nymphs were active mainly in May, while larvae were more active in July and adults in October and November. Overall, the seasonal tick activity in the environment and their abundance on the hosts peaked simultaneously, although the peak activity of questing *H. longicornis* adults and *I. scapularis* nymphs preceded their peak infestation on the hosts (Fig. 5). However, although adults of *A. americanum* were present at high densities in the environment from May to July (Supplementary Table S5), we found a single adult feeding on a raccoon in September.

## 4. Discussion

We captured and processed a variety of small and medium-sized mammal species over five months in suburban New Jersey, identified eight of the 11 species of Ixodida known to occur in the state (Occi et al., 2019) and detected *I. marxi* for the first time. The presence of *I. marxi* is not unexpected since this species was known to occur on squirrels in all neighboring states (Occi et al., 2019). The areas we sampled are human-modified environments used for recreational, agricultural, and gardening purposes all approximately 1 km apart inside a large state university campus. The proximity and variety of small forest fragments and ecotone areas supports a high diversity of mammals and provides the invasive *H. longicornis* and native ticks the opportunity to find and feed on a broad spectrum of hosts. Importantly, we found *H. longicornis* to be the numerically dominant tick species both on mammals and in the environment.

*Haemaphysalis longicornis* has a relatively wide host range in its native and invaded areas (Zhao et al., 2020), including mammals from the orders Rodentia, Eulipotyphla, Lagomorpha, Didelphimorphia, and Carnivora analyzed in our study. We detected *H. longicornis* feeding on eight different mammal species. Of note, we collected *H. longicornis* from one eastern chipmunk, a new host record (USDA, 2023), from one *Peromyscus* mouse, from 27% of the gray squirrels and 78% of groundhogs, showing it feeds on rodents more frequently than previously reported (Hoogstraal et al., 1968; Zheng et al., 2012; Tufts et al., 2019; Breuner et al., 2020). Interestingly, we found higher numbers of specimens on gray squirrels than on the other rodents even though they were all trapped in the same habitats teeming with *H. longicornis*. This difference could be explained by host preference, which can only be assessed with host-choice experiments.

Since the first detection in NJ in 2017, *H. longicornis* has been primarily recorded to feed on wildlife such as deer, Virginia opossum and raccoon, livestock (cows) and companion animals (dogs), and have been increasing in importance as a nuisance for humans (Beard et al., 2018; Bickerton and Toledo, 2020; Wormser et al., 2020; Thompson et al., 2021; Tufts et al., 2021; White et al., 2021). Therefore, it is not surprising that we found Virginia opossums and raccoons to be highly parasitized by *H. longicornis*. However, we observed a higher abundance of *H. longicornis* on striped skunks than other recent studies (Tufts et al., 2021; White et al., 2021). Although there are relatively few records of tick infestations in skunks, the main species reported were *A. americanum*, *D. variabilis*, *I. cookei* and *I. scapularis* (Zimmerman et al., 1988; Fish and Dowler, 1989; Durden and Richardson, 2003; Cohen et al., 2010). Our study adds robust information that *H. longicornis* is now a major tick species feeding on striped skunks in the US.

We observed a similar prevalence and host range for *H. longicornis* and *A. americanum* with the exception that *A. americanum* in our study was found only occasionally on Virginia opossums, a pattern others have also found (Childs and Paddock, 2003), while all stages of *H. longicornis* and immature *I. scapularis* predominated on opossums. Unlike the other animals captured, opossum are marsupials with a relatively slow metabolism and low body temperature (McManus, 1974), which may affect the tick feeding duration (Pollock et al., 2015). It appears that *H. longicornis* and *I. scapularis* have adapted to these conditions. The use of Virginia opossum by *I. scapularis* has been noted by others (Magnarelli et al., 1984; Fish and Dowler, 1989), and our study adds to the growing evidence that Virginia opossum are also important host for *H. longicornis*.

While it was not surprising to find that immature stages of *I. scapularis* predominated on small mammals, it is noteworthy that we observed the greatest abundance of *I. scapularis* nymphs on eastern chipmunks, with 36 of the 37 individuals sampled having immature stages of this tick species. Our findings of the importance of chipmunks as *I. scapularis* hosts support those of others (Schmidt et al., 1999; Sidge et al., 2021). Eastern chipmunks are diurnal animals, most active during mid-morning and mid-afternoon, that may spend less time grooming than *Peromyscus* mice, a well-known self-cleaning activity that can remove significant numbers of ticks (Yahner, 1978; Shaw et al., 2003). In addition, and in contrast to other authors (Shaw et al., 2003; Sidge et al., 2021), we collected a similar abundance of larvae of *I. scapularis* from *Peromyscus* sp. mice and eastern chipmunks. Our findings support the importance of eastern chipmunks as reservoir hosts of the Lyme spirochete *Borrelia burgdorferi* (McLean et al., 1993; Schmidt et al., 1999), especially since they are known to live longer than *Peromyscus* mice (Tryon and Snyder, 1973; Aguilar, 2011).

House mice and short-tailed shrews have low exposure to ticks because they stay primarily indoors and in burrows, respectively (Telford III et al., 1990; Mitsainas et al., 2016), but we found that both mammal species were commonly infested with immature *I. scapularis*, indicating they can host important disease vectors in the human modified areas we surveyed (Brillhart et al., 1994). While short-tailed shrews are insectivores in the order Eulipotyphla that do not have external ears, one of the preferred attachment sites for ticks (Smart and Caccamise, 1988), they are competent and important hosts of *Borrelia burgdorferi*, *Babesia microti,* and *Anaplasma phagocytophilum*, important zoonotic pathogens in the Northeastern USA (Telford III et al., 1990; LoGiudice et al., 2003; Hersh et al., 2012; Keesing et al., 2012). In contrast, the role of free-ranging house mice as pathogen reservoirs is unclear (Russart et al., 2014), probably because they are not commonly captured and tested for the presence of tick-borne pathogens.

Meadow voles hosted low numbers of ticks, but harbored the highest abundance of immature stages of *D. variabilis* as previously described (Smart and Caccamise, 1988; Kollars et al., 2000). Meadow voles are burrowing rodents that are active day and night, and in New Jersey occupy mostly grassy habitats rather than wooded areas where *I. scapularis* and *Peromyscus* mice are usually observed (Brooks and Webster, 1981; Smart and Caccamise, 1988). We also collected *D. variabilis* adults from larger mammals, such as skunks, opossums, groundhogs and especially raccoons, as reported by others (Kollars et al., 2000; Cohen et al., 2010). *Dermacentor variabilis* is a competent vector of *R. rickettsia*, the bacterium that causes Rocky Mountain spotted fever (Azad and Beard, 1998), *Francisella tularensis*, the agent of tularemia (Reese et al., 2011), and Powassan virus (POWV) under laboratory conditions (Sharma et al., 2021). Furthermore, Powassan virus (POWV) DNA was recently detected in a pool of *D. variabilis* females collected from the field in the state of New York (Hart et al., 2023). POWV is a pathogenic neurotropic tick-borne flavivirus that in the USA uses mainly *Ixodes* ticks (*I. scapularis* and *I. marxi*) and rodents as vectors and reservoirs, respectively (Hassett and Thangamani, 2021). However, those recent findings involving *D. variabilis* suggest that *Peromyscus* mice and meadow voles could be the important hosts of the immature ticks of this species in our study area. Voles have been suggested as potential reservoirs of POWV in the USA (Hassett and Thangamani, 2021), although one study found no seropositivity for POWV in meadow voles sampled in areas where 3% to 4% of *Peromyscus leucopus* had evidence of viral exposure (Ebel, 2010). Since this virus is an emerging zoonotic threat (Hassett and Thangamani, 2021), the enzootic cycle involving these small rodents and *D. variabilis* should be further investigated.

Raccoons harbored the greatest tick richness (6 out of 9 tick species) and were parasitized by the three stages of all tick species except for *D. variabilis*. In addition to *H. longicornis, A. americanum, I. scapularis* and *D. variabilis*, we collected *I. cookei* and *I. texanus* from raccoons. *Ixodes texanus* usually parasitize raccoons, but also feeds on skunks and opossums (Fish and Dowler, 1989; Pung et al., 1994; Kollars and Oliver, 2003; Cohen et al., 2010). Similarly, we also collected *I. cookei* from other mammals, especially skunks (Durden and Richardson, 2003; Cohen et al., 2010). Both *I. texanus* and *I. cookei* are nidicolous that when off-host remain in burrows used by their hosts (Anderson and Magnarelli, 2008; Ko, 1972), establishing a narrow host selection pattern that can minimize health risks for humans and animals. Even so, it should be noted that *I. texanus* infected with *B. burgdorferi* and *Babesia lotori* have been found in the eastern USA (Anderson et al., 1981; Ouellette et al., 1997). In addition, *I. cookei* is a known vector of POWV and groundhogs are considered the main reservoir for this virus in northeastern USA (Ebel, 2010), although the role of other hosts, such as skunks, is under investigation (Kemenesi and Bányai, 2019).

Our results indicate that heavier mammals, such as raccoons, opossums, and skunks, harbor the highest tick abundances, especially of adult *H. longicornis, A. americanum* and *D. variabilis,* emphasizing the importance of these hosts in the maintenance and spread of these tick species. Previous studies have shown a positive linear association between animal body weight and number of ticks, although other factors such as skin thickness, behavioral features discussed above, or immune response against tick bites are also important determinants of host suitability (Esser et al., 2016; Mysterud et al., 2021). For example, the low tick infestation observed in groundhogs could be because their thick skin hinders tick attachment, as suggested in a study with badgers (Mysterud et al., 2021). Others have found sex-biased parasitism due to the difference in body weight between the sexes (Kiffner et al., 2011; Krasnov et al., 2012), however, in our study while females tended to have less ticks, the results were not statistically significant.

The tick community composition and seasonal dynamics we observed in the environment followed expected patterns (González et al., 2023), and was characterized by high numbers of *H. longicornis*, followed by *A. americanum*, *I. scapularis* and *D. variabilis*. These patterns also reflect the seasonality of tick abundance on the captured animals for most tick species and stages. Our mammal surveillance revealed the presence of nidicolous tick stages and species, which are rarely collected from the environment, such as the immature *D. variabilis*, and all stages of *I. cookei*, *I. dentatus*, *I. marxi* and *I. texanus*. The fact that we collected virtually no adults of *A. americanum* from the mammals we trapped despite high environmental densities between May and July is likely because we did not sample its main host, the white-tailed deer (Schulze et al., 1984). Although sampling effort is critical, tick surveillance from animals is not as affected by climatic variables and vegetation structure as sampling host-seeking ticks (Estrada-Peña et al., 2013). Such investigations inform about tick community composition and tick ecology by uncovering new records of local vector species (e.g., *I. marxi* in New Jersey) and by determining patterns of host use for different tick species for all stages.

Our results showing seasonal tick dynamics on hosts and on the environment is crucial for guiding targeted control efforts and informing public and animal health recommendations on tick avoidance during peak activity periods. We showed that the newly invasive *H. longicornis* is now the most abundant tick species on mammals and in the environment in suburban areas within the NJ urban belt only four years after its first detection in New Jersey. We also found *H. longicornis* feeding on a diverse range of small and medium-sized mammals, many of which are known reservoirs of viruses, bacteria, and protozoa of medical and veterinary importance. Therefore, future studies should test *H. longicornis* collected from the environment and on their vertebrate hosts this species for a broad array of pathogens to determine the role of this vector species in enzootic and zoonotic disease cycles as they expand their territory in the US.

## Acknowledgements

We thank the Centers for Disease Control and Prevention (CDC) for providing the Sherman and medium Tomahawk traps used in this study and Dr. Lars Eisen for feedback on an earlier version of the manuscript. We are also indebted to the many graduate and undergraduate students, faculty and staff that gave us a hand and we needed it. Special thanks to Zoe Narvaez, Dr. Andrea Egizi, Nicole Wagner, and Dr. Jim Occi for assistance in the field. We thank Richard G. Robbins for his help in the identification of some tick species. We thank Dr. David Reimer for helping us develop the IACUC protocol, Clinton Burgher for providing us the keys to get into the farming areas, Rutgers Gardens people for facilitating our mammal trapping and Jacob Nava from Fish and Wildlife for guidance.

## Funding

This study was supported by the Cooperative Agreement Number 1U01CK000509 between the Centers for Disease Control and Prevention (CDC) and the Northeast Center of Excellence in Vector Borne Diseases (sub-contract to DMF). The contents of this work are solely the responsibility of the authors and do not necessarily represent the official views of the Centers for Disease Control and Prevention or the Department of Health and Human Services.

## CRediT authorship contribution statement

**Francisco C. Ferreira and Julia González:** Conceptualization, Methodology, Formal analysis, Investigation, Data curation, Writing – original draft, Visualization. **Matthew T. Milholland:** Methodology, Investigation, Writing - Review & Editing. **Grayson A. Tung:** Methodology, Investigation, Writing - Review & Editing. **Dina M. Fonseca:** Conceptualization, Methodology, Resources, Writing - Review & Editing, Supervision, Project Administration, Funding Acquisition.

## ORCID

Francisco C. Ferreira https://orcid.org/0000-0002-2034-4121

Julia González https://orcid.org/0000-0003-1517-4559

Matthew T. Milholland https://orcid.org/0000-0001-9398-7243

Dina M. Fonseca https://orcid.org/0000-0003-4726-7100

---

**Supplementary Table S1.** Tick infestation prevalence (number of infested individuals/total individuals sampled; also shown in percentage) and tick abundance (mean ticks per host species sampled) for each tick species collected in New Brunswick NJ, USA, 2021. The range of ticks (minimum – maximum) is also shown. *Haemaphysalis leporispalustris* and *Ixodes marxi* were not included because one and two specimens of each were collected from an eastern cottontail rabbit and from two eastern gray squirrels, respectively.

**Supplementary Table S1.** Tick infestation prevalence (number of infested individuals/total individuals sampled; also shown in percentage) and tick abundance (mean ticks per host species sampled) for each tick species collected in New Brunswick NJ, USA, 2021.
|  |  | house<br>mouse | short-tailed<br>shrew | *Peromyscus*<br>mouse | meadow<br>vole | eastern<br>chipmunk | gray<br>squirrel | eastern<br>cottontail | striped<br>skunk | Virginia<br>opossum | groundhog | raccoon | total |
| --- | --- | --- | --- | --- | --- | --- | --- | --- | --- | --- | --- | --- | --- |
| ***Amblyomma***<br>***americanum*** | Prevalence (%) | 0 | 0 | 1/78<br>(1.3) | 1/29<br>(3.4) | 0 | 26/37<br>(70.3) | 1/1<br>(100) | 12/13<br>(92.3) | 3/11<br>(27.3) | 7/9<br>(77.8) | 34/37<br>(91.9) | **85/273**<br>**(31.1)** |
|  | Abundance |  |  | 0.01 | 0.03 |  | 2.70 | 7.00 | 44.69 | 0.82 | 2.00 | 33.16 | **7.12** |
|  | Range of ticks |  |  | 0-1 | 0-1 |  | 0-16 | 7-7 | 0-134 | 0-4 | 0-5 | 0-190 | **0-190** |
| ***Dermacentor***<br>***variabilis*** | Prevalence (%) | 0 | 0 | 21/78<br>(26.9) | 13/29<br>(44.8) | 1/37<br>(2.7) | 0 | 0 | 3/13<br>(23.1) | 5/11<br>(45.5) | 3/9<br>(33.3) | 16/37<br>(43.2) | **62/273**<br>**(22.8)** |
|  | Abundance |  |  | 0.78 | 1.86 | 0.03 |  |  | 0.77 | 0.91 | 0.67 | 4.62 | **1.15** |
|  | Range of ticks |  |  | 0-10 | 0-12 | 0-1 |  |  | 0-7 | 0-4 | 0-3 | 0-37 | **0-37** |
| ***Haemaphysalis***<br>***longicornis*** | Prevalence (%) | 0 | 0 | 1/78<br>(1.3) | 0 | 1/37<br>(2.7) | 10/37<br>(27.0) | 1/1<br>(100) | 12/13<br>(92.3) | 10/11<br>(90.9) | 7/9<br>(77.8) | 34/37<br>(91.9) | **76/273**<br>**(27.8)** |
|  | Abundance |  |  | 0.01 |  | 0.03 | 1.19 | 6.00 | 34.38 | 65.91 | 1.89 | 52.81 | **11.70** |
|  | Range of ticks |  |  | 0-1 |  | 0-1 | 0-29 | 6-6 | 0-133 | 0-301 | 0-8 | 0-333 | **0-333** |
| ***Ixodes***<br>***cookei*** | Prevalence (%) | 0 | 0 | 0 | 0 | 0 | 0 | 0 | 9/13<br>(69.2) | 1/11<br>(9.1) | 2/9<br>(22.2) | 12/37<br>(32.4) | **24/273**<br>**(8.8)** |
|  | Abundance |  |  |  |  |  |  |  | 7.31 | 0.09 | 0.44 | 1.30 | **0.52** |
|  | Range of ticks |  |  |  |  |  |  |  | 0-45 | 0-1 | 0-3 | 0-15 | **0-45** |
| ***Ixodes***<br>***dentatus*** | Prevalence (%) | 0 | 0 | 0 | 0 | 0 | 0 | 1/1<br>(100) | 0 | 0 | 0 | 0 | **1/273**<br>**(0.37)** |
|  | Abundance |  |  |  |  |  |  | 9.00 |  |  |  |  | **0.03** |
|  | Range of ticks |  |  |  |  |  |  | 9-9 |  |  |  |  | **0-9** |
| ***Ixodes***<br>***scapularis*** | Prevalence (%) | 11/16<br>(68.7) | 4/5<br>(80.0) | 63/78<br>(80.8) | 12/29<br>(41.4) | 36/37<br>(97.3) | 27/37<br>(73.0) | 0 | 12/13<br>(92.3) | 11/11<br>(100) | 9/9<br>(100) | 27/37<br>(73.0) | **212/273**<br>**(77.6)** |
|  | Abundance | 2.13 | 8.20 | 12.65 | 0.86 | 18.19 | 8.54 |  | 15.85 | 51.27 | 7.67 | 4.32 | **11.27** |
|  | Range of ticks | 0-12 | 0-29 | 0-137 | 0-6 | 0-101 | 0-93 |  | 0-100 | 3-172 | 1-35 | 0-51 | **0-172** |
| ***Ixodes***<br>***texanus*** | Prevalence (%) | 0 | 0 | 0 | 0 | 0 | 0 | 0 | 0 | 0 | 0 | 21/37<br>(56.8) | **21/273**<br>**(7.7)** |
|  | Abundance |  |  |  |  |  |  |  |  |  |  | 8.76 | **1.19** |
|  | Range of ticks |  |  |  |  |  |  |  |  |  |  | 0-98 | **0-98** |
| **total ticks** | Prevalence (%) | **11/16**<br>**(68.7)** | **4/5**<br>**(80.0)** | **68/78**<br>**(87.2)** | **20/29**<br>**(69.0)** | **36/37**<br>**(97.3)** | **34/37**<br>**(91.9)** | **1/1**<br>**(100)** | **12/13**<br>**(92.3)** | **11/11**<br>**(100)** | **9/9**<br>**(100)** | **37/37**<br>**(100)** | **243/273**<br>**(89.0)** |
|  | Abundance | **2.13** | **8.20** | **13.46** | **2.76** | **18.24** | **12.46** | **23** | **103.0** | **119.0** | **12.67** | **104.90** | **33.0** |
|  | Range of ticks | **0-12** | **0-29** | **0-137** | **0-12** | **0-101** | **0-102** | **23-23** | **0-257** | **22-304** | **2-41** | **1-562** | **0-562** |

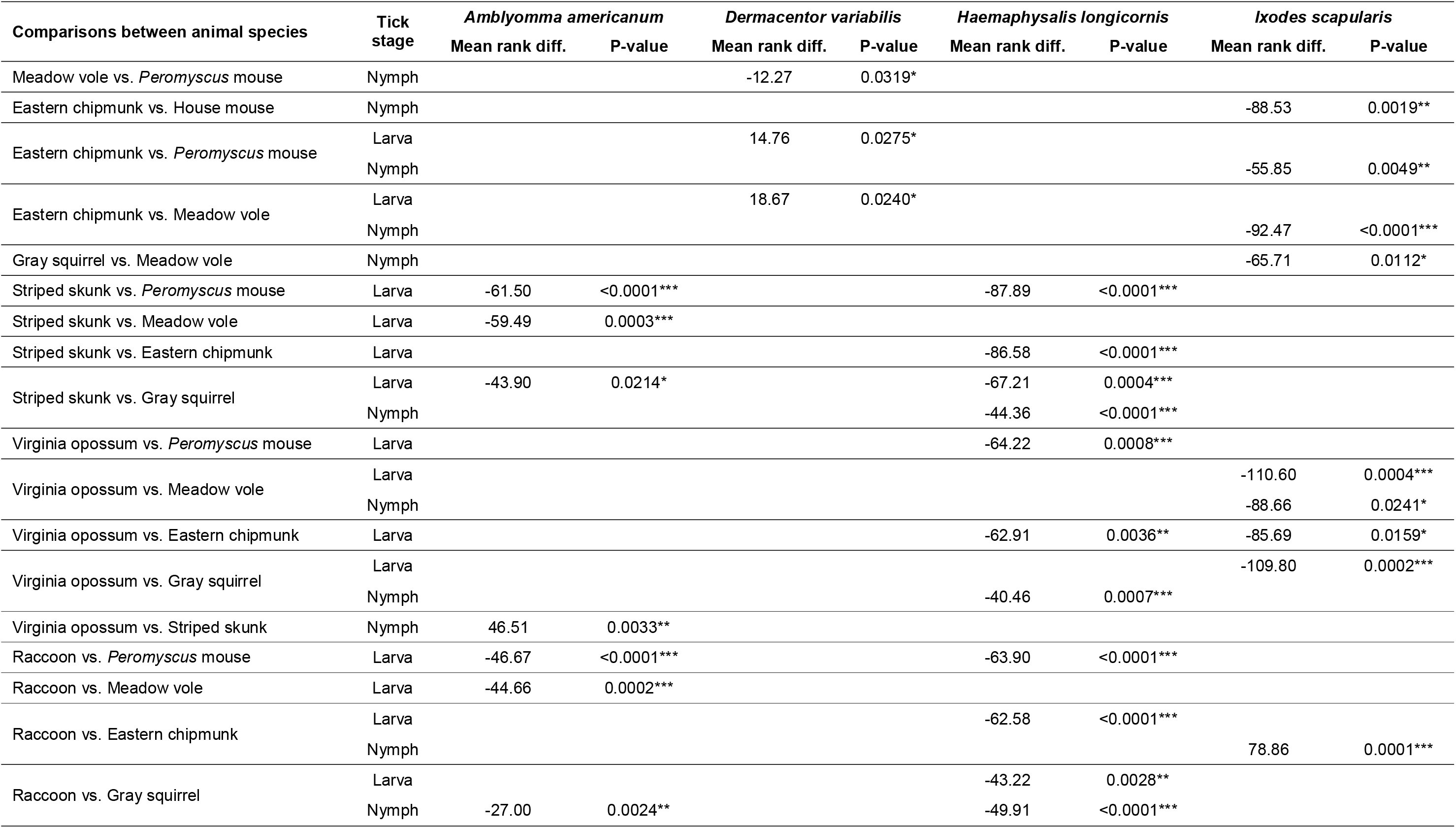
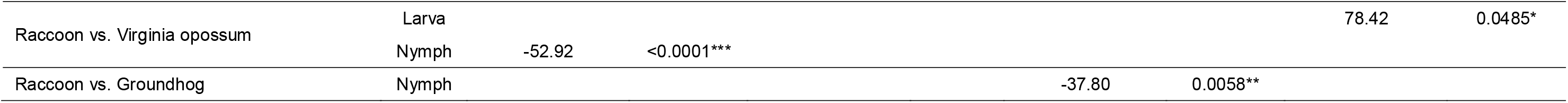
**Supplementary Table S2.** Significant results obtained by Dunn’s multiple comparisons tests between host species captured. Previous Kruskal-Wallis tests showed significant differences (P < 0.05) in the number of larvae and nymphs collected between hosts for the main four tick species. Results of adult ticks collected are not included because the Kruskal-Wallis tests showed no significant variation between medians.

**Supplementary Table S3.** Spearman correlation between abundance of the four main tick species (*A. americanum*, *D. variabilis*, *H. longicornis* and *I. scapularis*) and the mean weight of their hosts analyzed by tick life stage. Significant relationships were then analyzed by simple linear regression. P < 0.05 (*), P < 0.01 (**) and P < 0.001 (***).

**Supplementary Table S3.** Spearman correlation between abundance of the four main tick species (*A.
| Tick species | Tick stage | Spearman correlation vs weight of animals |  |  | Simple linear regression vs weight of animals |  |  |  |  |
| --- | --- | --- | --- | --- | --- | --- | --- | --- | --- |
|  |  | rs | 95%<br>Confidence Interval | P value | Slope | Std.<br>Error | R<sup>2</sup> | F<sub>1,266</sub> | P value |
| *Amblyomma americanum* | Larvae | 0.387 | 0.277 to 0.487 | <0.0001*** | 0.0049 | 0.0008 | 0.129 | 39.26 | <0.0001*** |
|  | Nymphs | 0.678 | 0.605 to 0.739 | <0.0001*** | 0.0015 | 0.0002 | 0.200 | 66.50 | <0.0001*** |
|  | Adults | 0.102 | -0.021 to 0.223 | 0.094 |  |  |  |  |  |
| *Dermacentor variabilis* | Larvae | -0.232 | -0.345 to -0.112 | 0.0001*** | -0.0001 | 4.97 x10<sup>-5</sup> | 0.016 | 4.398 | 0.037* |
|  | Nymphs | -0.139 | -0.258 to -0.016 | 0.022* | -3.83 x10<sup>-5</sup> | 1.96 x10<sup>-5</sup> | 0.014 | 3.823 | 0.052 |
|  | Adults | 0.469 | 0.367 to 0.560 | <0.0001*** | 0.0011 | 0.0001 | 0.251 | 89.06 | <0.0001*** |
| *Haemaphysalis longicornis* | Larvae | 0.520 | 0.424 to 0.605 | <0.0001*** | 0.0083 | 0.0015 | 0.100 | 29.69 | <0.0001*** |
|  | Nymphs | 0.661 | 0.585 to 0.725 | <0.0001*** | 0.0012 | 0.0002 | 0.174 | 56.19 | <0.0001*** |
|  | Adults | 0.295 | 0.178 to 0.403 | <0.0001*** | 0.0005 | 9.98 x10<sup>-5</sup> | 0.095 | 27.85 | <0.0001*** |
| *Ixodes scapularis* | Larvae | 0.002 | -0.121 to 0.126 | 0.966 |  |  |  |  |  |
|  | Nymphs | 0.120 | -0.003 to 0.240 | 0.049* | -0.0003 | 0.0002 | 0.009 | 2.354 | 0.126 |
|  | Adults | 0.098 | -0.025 to 0.219 | 0.108 |  |  |  |  |  |

**Supplementary Table S4.** Number of ticks collected by flagging (adults; nymphs; larvae) in New Brunswick NJ, USA, 2021.

| Month | samplings | *Amblyomma americanum* | *Dermacentor variabilis* | *Haemaphysalis leporispalustris* | *Haemaphysalis longicornis* | *Ixodes scapularis* | Total ticks |
| --- | --- | --- | --- | --- | --- | --- | --- |
| May | 1 | 13; 24; 0 | 0; 0; 0 | 0; 0; 0 | 8; 97; 463 | 3; 35; 1 | 644 |
| June | 1 | 33; 9; 0 | 0; 0; 0 | 0; 0; 0 | 31; 113; 25 | 0; 7; 0 | 218 |
| July | 4 | 50; 93; 175 | 3; 0; 0 | 0; 1; 0 | 132; 327; 13 | 0; 28; 1800 | 2622 |
| August | 2 | 0; 14; 2818 | 1; 0; 0 | 0; 0; 1 | 31; 46; 1222 | 0; 3; 402 | 4538 |
| September | 1 | 0; 2; 625 | 0; 0; 0 | 0; 0; 0 | 4; 9; 1358 | 0; 4; 35 | 2037 |
| October | 1 | 0; 0; 0 | 0; 0; 0 | 0; 0; 0 | 0; 0; 698 | 17; 4; 4 | 723 |
| November | 1 | 0; 0; 0 | 0; 0; 0 | 0; 0; 0 | 0; 0; 28 | 26; 0; 0 | 54 |
| **total** | **11** | **96; 142; 3618** | **4; 0; 0** | **0; 1; 1** | **206; 592; 3807** | **46; 81; 2242** | **10836** |

**Supplementary Table S5.** Number of ticks collected per square meter at three different sites in New Brunswick NJ, USA, 2021. Four *Dermacentor variabilis* adults, one nymphal and one larval *Haemaphysalis leporispalustris* were collected but are not included in the table. GF = Goat Farm; GR = Rutgers Gardens; UI = University Inn.

**Supplementary Table S5.** Number of ticks collected per square meter at three different sites in New Brunswick NJ, USA, 2021.
| month | samplings | *Amblyomma americanum* |  |  |  | *Haemaphysalis longicornis* |  |  |  | *Ixodes scapularis* |  |  |  | Total ticks/m<sup>2</sup> |  |  |  |
| --- | --- | --- | --- | --- | --- | --- | --- | --- | --- | --- | --- | --- | --- | --- | --- | --- | --- |
|  |  | All sites | GF | RG | UI | All sites | GF | RG | UI | All sites | GF | RG | UI | All sites | GF | RG | UI |
| May | 1 | **0.25** | 0.3 | 0.4 | 0.0 | **3.79** | 4.9 | 0.2 | 6.3 | **0.26** | 0.6 | 0.2 | 0.0 | **4.3** | 5.8 | 0.7 | 6.3 |
| Jun | 1 | **0.23** | 0.1 | 0.6 | 0.0 | **0.94** | 0.5 | 0.6 | 1.9 | **0.04** | 0.1 | 0.0 | 0.0 | **1.2** | 0.7 | 1.2 | 2.0 |
| Jul | 4 | **0.41** | 0.2 | 0.8 | 0.3 | **0.61** | 0.4 | 0.2 | 1.4 | **2.37** | 1.5 | 1.6 | 4.2 | **3.4** | 2.1 | 2.6 | 5.8 |
| Aug | 2 | **7.26** | 6.8 | 5.6 | 9.5 | **3.33** | 1.5 | 0.2 | 8.5 | **1.04** | 1.2 | 0.6 | 1.4 | **11.6** | 9.5 | 6.4 | 19.4 |
| Sep | 1 | **4.48** | 1.2 | 0.9 | 13.1 | **9.79** | 7.6 | 2.5 | 21.7 | **0.28** | 0.3 | 0.1 | 0.5 | **14.6** | 9.1 | 3.5 | 35.2 |
| Oct | 1 | **0** | 0.0 | 0.0 | 0.0 | **4.36** | 6.6 | 2.7 | 3.9 | **0.16** | 0.1 | 0.3 | 0.1 | **4.5** | 6.6 | 3.0 | 4.0 |
| Nov | 1 | **0** | 0.0 | 0.0 | 0.0 | **0.19** | 0.0 | 0.6 | 0.0 | **0.17** | 0.1 | 0.4 | 0.0 | **0.4** | 0.1 | 0.9 | 0.0 |
| **total** | **11** | **2.0** | **1.3** | **1.7** | **3.0** | **2.4** | **1.9** | **0.6** | **4.7** | **1.2** | **0.9** | **0.8** | **2.0** | **5.6** | **4.2** | **3.2** | **9.7** |

**Supplementary Material Fig. S1.** Relationship between tick abundance (left-Y axis) and mean weight of the host species captured (right-Y axis). The four most prevalen tick species, *Haemaphysalis longicornis*, *Amblyomma americanum*, *Dermacentor variabilis,* and *Ixodes scapularis,* are included. Refer to the legend for the colors representing each species. The graphs refer to each life stage: A) adults, B) nymphs and C) larvae. If zero ticks were collected, they are not represented.

## Notes

### Competing Interest Statement

The authors have declared no competing interest.

## References

Aguilar, S., 2011. *Peromyscus leucopus* (white-footed mouse) [WWW Document]. Animal Diversity Web. URL https://animaldiversity.org/accounts/Peromyscus_leucopus/ (accessed 12.23.22).

Altschul, S.F., Gish, W., Miller, W., Myers, E.W., Lipman, D.J., 1990. Basic local alignment search tool. J Mol Biol 215, 403–410. https://doi.org/10.1016/S0022-2836(05)80360-2

Anderson, J.F., Magnarelli, L.A., 2008. Biology of Ticks. Infectious Disease Clinics of North America 22, 195– 215. https://doi.org/10.1016/j.idc.2007.12.006

Anderson, J.F., Magnarelli, L.A., Sulzer, A.J., 1981. Raccoon babesiosis in Connecticut, USA: *Babesia lotori* sp. n. The Journal of Parasitology 67, 417–425. https://doi.org/10.2307/3280566

Azad, A.F., Beard, C.B., 1998. Rickettsial pathogens and their arthropod vectors. Emerg Infect Dis 4, 179–186. https://doi.org/10.3201/eid0402.980205

Bai, Q., Liu, G., Liu, D., Ren, J., Li, X., 2002. Isolation and preliminary characterization of a large *Babesia* sp. from sheep and goats in the eastern part of Gansu Province, China. Parasitol Res 88, S16–21. https://doi.org/10.1007/s00436-001-0563-6

Beard, C.B., Occi, J., Bonilla, D.L., Egizi, A.M., Fonseca, D.M., Mertins, J.W., Backenson, B.P., Bajwa, W.I., Barbarin, A.M., Bertone, M.A., Brown, J., Connally, N.P., Connell, N.D., Eisen, R.J., Falco, R.C., James, A.M., Krell, R.K., Lahmers, K., Lewis, N., Little, S.E., Neault, M., Pérez de León, A.A., Randall, A.R., Ruder, M.G., Saleh, M.N., Schappach, B.L., Schroeder, B.A., Seraphin, L.L., Wehtje, M., Wormser, G.P., Yabsley, M.J., Halperin, W., 2018. Multistate Infestation with the Exotic Disease–Vector Tick *Haemaphysalis longicornis* — United States, August 2017–September 2018. MMWR Morb. Mortal. Wkly. Rep. 67, 1310–1313. https://doi.org/10.15585/mmwr.mm6747a3

Bickerton, M., McSorley, K., Toledo, A., 2021. A life stage-targeted acaricide application approach for the control of *Haemaphysalis longicornis*. Ticks and Tick-borne Diseases 12, 101581. https://doi.org/10.1016/j.ttbdis.2020.101581

Bickerton, M., Toledo, A., 2020. Multiple pruritic tick bites by Asian Longhorned tick larvae (*Haemaphysalis longicornis*). International Journal of Acarology 46, 373–376. https://doi.org/10.1080/01647954.2020.1805004

Bowman, D., Little, S.E., Lorentzen, L., Shields, J., Sullivan, M.P., Carlin, E.P., 2009. Prevalence and geographic distribution of *Dirofilaria immitis, Borrelia burgdorferi, Ehrlichia canis*, and *Anaplasma phagocytophilum* in dogs in the United States: results of a national clinic-based serologic survey. Vet Parasitol 160, 138–148. https://doi.org/10.1016/j.vetpar.2008.10.093

Bozek, C.K., Prange, S., Gehrt, S.D., 2007. The influence of anthropogenic resources on multi-scale habitat selection by raccoons. Urban Ecosyst 10, 413–425. https://doi.org/10.1007/s11252-007-0033-8

Breuner, N.E., Ford, S.L., Hojgaard, A., Osikowicz, L.M., Parise, C.M., Rosales Rizzo, M.F., Bai, Y., Levin, M.L., Eisen, R.J., Eisen, L., 2020. Failure of the Asian longhorned tick, *Haemaphysalis longicornis*, to serve as an experimental vector of the Lyme disease spirochete, *Borrelia burgdorferi* sensu stricto. Ticks and Tick-borne Diseases 11, 101311. https://doi.org/10.1016/j.ttbdis.2019.101311

Brillhart, D.B., Fox, L.B., Upton, S.J., 1994. Ticks (Acari: Ixodidae) Collected from Small and Medium-Sized Kansas Mammals. Journal of Medical Entomology 31, 500–504. https://doi.org/10.1093/jmedent/31.3.500

Brooks, R.J., Webster, A.B., 1981. Social behavior and activity patterns of meadow voles in relation to seasonal change and snow cover. Eastern Pine and Meadow Vole Symposia. 64–71.

Brown, L.D., Cai, T.T., DasGupta, A., 2001. Interval Estimation for a Binomial Proportion. Statistical Science 16, 101–133. https://doi.org/10.1214/ss/1009213286

Bureau, U.C., 2021. Historical Population Density Data (1910-2020) [WWW Document]. Census.gov. URL https://www.census.gov/data/tables/time-series/dec/density-data-text.html (accessed 10.24.22).

Bush, A.O., Lafferty, K.D., Lotz, J.M., Shostak, A.W., 1997. Parasitology Meets Ecology on Its Own Terms: Margolis et al. Revisited. The Journal of Parasitology 83, 575. https://doi.org/10.2307/3284227

Casel, M.A., Park, S.J., Choi, Y.K., 2021. Severe fever with thrombocytopenia syndrome virus: emerging novel phlebovirus and their control strategy. Exp Mol Med 53, 713–722. https://doi.org/10.1038/s12276-021-00610-1

Childs, J.E., Paddock, C.D., 2003. The ascendancy of *Amblyomma americanum* as a vector of pathogens affecting humans in the United States. Annu. Rev. Entomol. 48, 307–337. https://doi.org/10.1146/annurev.ento.48.091801.112728

Chitimia, L., Lin, R.-Q., Cosoroaba, I., Wu, X.-Y., Song, H.-Q., Yuan, Z.-G., Zhu, X.-Q., 2010. Genetic characterization of ticks from southwestern Romania by sequences of mitochondrial cox1 and nad5 genes. Exp Appl Acarol 52, 305–311. https://doi.org/10.1007/s10493-010-9365-9

Cohen, S.B., Freye, J.D., Dunlap, B.G., Dunn, J.R., Jones, T.F., Moncayo, A.C., 2010. Host Associations of *Dermacentor*, *Amblyomma*, and *Ixodes* (Acari: Ixodidae) Ticks in Tennessee. J. Med. Entomol. 47, 415–420. https://doi.org/10.1603/ME09065

Cumbie, A.N., Trimble, R.N., Eastwood, G., 2022. Pathogen Spillover to an Invasive Tick Species: First Detection of Bourbon Virus in *Haemaphysalis longicornis* in the United States. Pathogens 11, 454. https://doi.org/10.3390/pathogens11040454

Dinkel, K.D., Herndon, D.R., Noh, S.M., Lahmers, K.K., Todd, S.M., Ueti, M.W., Scoles, G.A., Mason, K.L., Fry, L.M., 2021. A U.S. isolate of *Theileria orientalis*, Ikeda genotype, is transmitted to cattle by the invasive Asian longhorned tick, Haemaphysalis longicornis. Parasites Vectors 14, 157. https://doi.org/10.1186/s13071-021-04659-9

Durden, L.A., Keirans, J.E., 1996. Nymphs of the Genus *Ixodes* (Acari: Ixodidae) of the United States: Taxonomy, Identification Key, Distribution, Hosts, and Medical/veterinary Importance. Entomological Society of America.

Durden, L.A., Richardson, D.J., 2003. Ectoparasites of the Striped Skunk, *Mephitis mephitis*, in Connecticut, U.S.A. Compar. Parasitol. 70, 42–45. https://doi.org/10.1654/1525-2647(2003)070[0042:EOTSSM]2.0.CO;2

Ebel, G.D., 2010. Update on Powassan Virus: Emergence of a North American Tick-Borne Flavivirus. Annu. Rev. Entomol. 55, 95–110. https://doi.org/10.1146/annurev-ento-112408-085446

Egizi, A., Bulaga-Seraphin, L., Alt, E., Bajwa, W.I., Bernick, J., Bickerton, M., Campbell, S.R., Connally, N., Doi, K., Falco, R.C., Gaines, D.N., Greay, T.L., Harper, V.L., Heath, A.C.G., Jiang, J., Klein, T.A., Maestas, L., Mather, T.N., Occi, J.L., Oskam, C.L., Pendleton, J., Teator, M., Thompson, A.T., Tufts, D.M., Umemiya-Shirafuji, R., VanAcker, M.C., Yabsley, M.J., Fonseca, D.M., 2020. First glimpse into the origin and spread of the Asian longhorned tick, *Haemaphysalis longicornis*, in the United States. Zoonoses Public Health 67, 637–650. https://doi.org/10.1111/zph.12743

Egizi, A.M., Occi, J.L., Price, D.C., Fonseca, D.M., 2019a. Leveraging the Expertise of the New Jersey Mosquito Control Community to Jump Start Standardized Tick Surveillance. Insects 10, 219. https://doi.org/10.3390/insects10080219

Egizi, A.M., Robbins, R.G., Beati, L., Nava, S., Evans, C.R., Occi, J.L., Fonseca, D.M., 2019b. A pictorial key to differentiate the recently detected exotic Haemaphysalis longicornis Neumann, 1901 (Acari, Ixodidae) from native congeners in North America. ZK 818, 117–128. https://doi.org/10.3897/zookeys.818.30448

Eisen, R.J., Eisen, L., Ogden, N.H., Beard, C.B., 2016. Linkages of weather and climate with *Ixodes scapularis* and *Ixodes pacificus* (Acari: Ixodidae), enzootic transmission of *Borrelia burgdorferi*, and Lyme disease in North America. J. Med. Entomol. 53, 250–261. https://doi.org/10.1093/jme/tjv199

Esser, H.J., Foley, J.E., Bongers, F., Herre, E.A., Miller, M.J., Prins, H.H.T., Jansen, P.A., 2016. Host body size and the diversity of tick assemblages on Neotropical vertebrates. Int. J. for Parasitol Wildl. 5, 295– 304. https://doi.org/10.1016/j.ijppaw.2016.10.001

Estrada-Peña, A., Cevidanes, A., Sprong, H., Millán, J., 2021. Pitfalls in Tick and Tick-Borne Pathogens Research, Some Recommendations and a Call for Data Sharing. Pathogens 10, 712. https://doi.org/10.3390/pathogens10060712

Estrada-Peña, A., Gray, J.S., Kahl, O., Lane, R.S., Nijhof, A.M., 2013. Research on the ecology of ticks and tick-borne pathogens—methodological principles and caveats. Front. Cell. Infect. Microbiol. 3. https://doi.org/10.3389/fcimb.2013.00029

Fish, D., Dowler, R.C., 1989. Host associations of ticks (Acari: Ixodidae) parasitizing medium-sized mammals in a Lyme disease endemic area of southern New York. J. Med. Entomol. 26, 200–209. https://doi.org/10.1093/jmedent/26.3.200

Fleshman, A.C., Foster, E., Maes, S.E., Eisen, R.J., 2022. Reported county-level distribution of seven human pathogens detected in host-seeking *Ixodes scapularis* and *Ixodes pacificus* (Acari: Ixodidae) in the contiguous United States. J. Med. Entomol. 59, 1328–1335. https://doi.org/10.1093/jme/tjac049

Folmer, O., Black, M., Hoeh, W., Lutz, R., Vrijenhoek, R., 1994. DNA primers for amplification of mitochondrial cytochrome c oxidase subunit I from diverse metazoan invertebrates. Mol. Mar. Biol. Biotechnol. 3, 294–299.

Ginsberg, H.S., Hickling, G.J., Burke, R.L., Ogden, N.H., Beati, L., LeBrun, R.A., Arsnoe, I.M., Gerhold, R., Han, S., Jackson, K., Maestas, L., Moody, T., Pang, G., Ross, B., Rulison, E.L., Tsao, J.I., 2021. Why Lyme disease is common in the northern US, but rare in the south: The roles of host choice, host-seeking behavior, and tick density. PLoS Biol. 19, e3001066. https://doi.org/10.1371/journal.pbio.3001066

Goddard, J., Varela-Stokes, A.S., 2009. Role of the lone star tick, *Amblyomma americanum* (L.), in human and animal diseases. Vet. Parasitol. 160, 1–12. https://doi.org/10.1016/j.vetpar.2008.10.089

González, J., Fonseca, D.M., Toledo, A., 2023. Seasonal dynamics of tick species in the ecotone of parks and recreational areas in Middlesex County (New Jersey, USA). Insects 14, 258. https://doi.org/10.3390/insects14030258

Guan, G., Moreau, E., Liu, J., Hao, X., Ma, M., Luo, J., Chauvin, A., Yin, H., 2010. Babesia sp. BQ1 (Lintan): molecular evidence of experimental transmission to sheep by *Haemaphysalis qinghaiensis* and Haemaphysalis longicornis. Parasitol. Int. 59, 265–267. https://doi.org/10.1016/j.parint.2009.12.002

Hart, C., Hassett, E., Vogels, C.B.F., Shapley, D., Grubaugh, N.D., Thangamani, S., 2023. Powassan Virus Lineage I in field-collected *Dermacentor variabilis ticks*, New York, USA. Emerg. Infect. Dis. 29, 3. https://doi.org/10.3201/eid2902.220410

Hassett, E.M., Thangamani, S., 2021. Ecology of Powassan virus in the United States. Microorganisms 9, 2317. https://doi.org/10.3390/microorganisms9112317

Hersh, M.H., Tibbetts, M., Strauss, M., Ostfeld, R.S., Keesing, F., 2012. Reservoir competence of wildlife host species for *Babesia microti*. Emerg. Infect. Dis. 18, 1951–1957. https://doi.org/10.3201/eid1812.111392

Hoogstraal, H., Roberts, F.H.S., Kohls, G.M., Tipton, V.J., 1968. Review of *Haemaphysalis* (Kaiseriana) *longicornis* Neumann (Resurrected) of Australia, New Zealand, New Caledonia, Fiji, Japan, Korea, and Northestern China and USSR and its parthenogenetic and bisexual populations (Ixodoidea, Ixodidae). J. of Parasitol. 54, 1197–1213.

Hutcheson, K., 1970. A test for comparing diversities based on the Shannon formula. J. Theor. Biol. 29, 151– 154. https://doi.org/10.1016/0022-5193(70)90124-4

Johnson, B.J., Robson, M.G., Fonseca, D.M., 2015. Unexpected spatiotemporal abundance of infected *Culex restuans* suggest a greater role as a West Nile virus vector for this native species. Infect. Genet. Evol. 31, 40–47. https://doi.org/10.1016/j.meegid.2015.01.007

Keesing, F., Hersh, M.H., Tibbetts, M., McHenry, D.J., Duerr, S., Brunner, J., Killilea, M., LoGiudice, K., Schmidt, K.A., Ostfeld, R.S., 2012. Reservoir competence of vertebrate hosts for *Anaplasma phagocytophilum*. Emerg. Infect. Dis. 18, 2013–2016. https://doi.org/10.3201/eid1812.120919

Keirans, J.E., Durden, L.A., 1998. Illustrated key to nymphs of the tick genus *Amblyomma* (Acari: Ixodidae) found in the United States. J. Med. Entomol. 35, 489–495. https://doi.org/10.1093/jmedent/35.4.489

Kemenesi, G., Bányai, K., 2019. Tick-Borne flaviviruses, with a focus on Powassan virus. Clin. Microbiol. Rev. 32, e00106–17. https://doi.org/10.1128/CMR.00106-17

Kiffner, C., Vor, T., Hagedorn, P., Niedrig, M., Rühe, F., 2011. Factors affecting patterns of tick parasitism on forest rodents in tick-borne encephalitis risk areas, Germany. Parasitol. Res. 108, 323–335. https://doi.org/10.1007/s00436-010-2065-x

Ko, R.C., 1972. Biology of *Ixodes cookei* Packard (Ixodidae) of groundhogs (*Marmota monax* Erxleben). Can. J. Zool. 50, 433–436.

Kollars, T.M., Oliver, J.H., 2003. Host associations and seasonal occurrence of *Haemaphysalis leporispalustris*, *Ixodes brunneus*, *I. cookei*, *I. dentatus*, and *I. texanus* (Acari: Ixodidae) in southeastern Missouri. J Med. Entomol. 40, 103–107. https://doi.org/10.1603/0022-2585-40.1.103

Kollars, T.M., Oliver Jr, J.H., Masters, E.J., Kollars, P.G., Durden, L.A., 2000. Host utilization and seasonal occurrence of Dermacentor species (Acari: Ixodidae) in Missouri. Exper. Appl. Acarol. 24, 631–643.

Krasnov, B.R., Bordes, F., Khokhlova, I.S., Morand, S., 2012. Gender-biased parasitism in small mammals: patterns, mechanisms, consequences. Mammalia 76, 1–13. https://doi.org/10.1515/mammalia-2011-0108

Krasnov, B.R., Stanko, M., Morand, S., 2007. Host community structure and infestation by ixodid ticks: repeatability, dilution efect and ecological specialization. Oecologia 154, 185–194.

Linske, M.A., Williams, S.C., Stafford, K.C., Ortega, I.M., 2018. *Ixodes scapularis* (Acari: Ixodidae) reservoir host diversity and abundance impacts on dilution of *Borrelia burgdorferi* (Spirochaetales: Spirochaetaceae) in residential and woodland habitats in Connecticut, United States. J. Med. Entomol. 55, 681–690. https://doi.org/10.1093/jme/tjx237

LoGiudice, K., Ostfeld, R.S., Schmidt, K.A., Keesing, F., 2003. The ecology of infectious disease: effects of host diversity and community composition on Lyme disease risk. Proc. Natl. Acad. Sci. USA. 100, 567–571. https://doi.org/10.1073/pnas.0233733100

Madison-Antenucci, S., Kramer, L.D., Gebhardt, L.L., Kauffman, E., 2020. Emerging Tick-Borne Diseases. Clin. Microbiol. Rev. 33. https://doi.org/10.1128/CMR.00083-18

Magnarelli, L.A., Anderson, J.F., Burgdorfer, W., Chappell, W.A., 1984. Parasitism by *Ixodes dammini* (Acari: Ixodidae) and antibodies to Spirochetes in mammals at Lyme Disease Foci in Connecticut, USA. J. Med. Entomol. 21, 52–57. https://doi.org/10.1093/jmedent/21.1.52

Mahachi, K., Kontowicz, E., Anderson, B., Toepp, A.J., Lima, A.L., Larson, M., Wilson, G., Grinnage-Pulley, T., Bennett, C., Ozanne, M., Anderson, M., Fowler, H., Parrish, M., Saucier, J., Tyrrell, P., Palmer, Z., Buch, J., Chandrashekar, R., Scorza, B., Brown, G., Oleson, J.J., Petersen, C.A., 2020. Predominant risk factors for tick-borne co-infections in hunting dogs from the USA. Parasit. Vectors 13, 247. https://doi.org/10.1186/s13071-020-04118-x

Margolis, L., Esch, G.W., Holmes, J.C., Kuris, A.M., Schad, G.A., 1982. The use of ecological terms in parasitology (report of an ad hoc committee of the American Society of Parasitologists). J. Parasitol. 68, 131–133.

McLean, R.G., Ubico, S.R., Cooksey, L.M., 1993. Experimental infection of the eastern chipmunk (*Tamias striatus*) with the Lyme disease spirochete (*Borrelia burgdorferi*). J. Wildl. Dis. 29, 527–532.

McManus, J.J., 1974. Didelphis virginiana. Mammalian Species 40, 1–6. https://doi.org/10.2307/3503783

Mitsainas, G., Yigit, N., Kryštufek, B., Musser, G., Group, R.H., 2016. IUCN Red List of Threatened Species: *Mus musculus*. IUCN Red List of Threatened Species.

Mysterud, A., Hügli, C., Viljugrein, H., 2021. Tick infestation on medium–large-sized mammalian hosts: are all equally suitable to *Ixodes ricinus* adults? Parasit. Vectors 14, 254. https://doi.org/10.1186/s13071-021-04775-6

Nieto, N.C., Porter, W.T., Wachara, J.C., Lowrey, T.J., Martin, L., Motyka, P.J., Salkeld, D.J., 2018. Using citizen science to describe the prevalence and distribution of tick bite and exposure to tick-borne diseases in the United States. PLoS ONE 13, e0199644. https://doi.org/10.1371/journal.pone.0199644

Occi, J.L., Egizi, A.M., Robbins, R.G., Fonseca, D.M., 2019. Annotated List of the Hard Ticks (Acari: Ixodida: Ixodidae) of New Jersey. J. Med. Entomol.56, 589–598. https://doi.org/10.1093/jme/tjz010

Oksanen, J., Blanchet, F.G., Friendly, M., Kindt, R., Legendre, P., McGlinn, D., Minchin, P.R., O’Hara, R.B., Simpson, G.L., Solymos, P., 2020. Vegan: Community ecology package. Ordination methods, diversity analysis and other functions for community and vegetation ecologists. R package version 2.5 (2019). R Package Version. Available online: https://CRAN. R-project. org/package= vegan (accessed on 13 December 2021).

Ondrejicka, D.A., Morey, K.C., Hanner, R.H., 2017. DNA barcodes identify medically important tick species in Canada. Genome 60, 74–84. https://doi.org/10.1139/gen-2015-0179

Ouellette, J., Apperson, C.S., Howard, P., Evans, T.L., Levine, J.F., 1997. Tick-raccoon associations and the potencial for Lyme disease spirochete transmission in the coastal plain of North Carolina. J. Wildl. Dis. 33, 28–39.

Pollock, N.B., Gawne, E., Taylor, E.N., 2015. Effects of temperature on feeding duration, success, and efficiency of larval western black-legged ticks (Acari: Ixodidae) on western fence lizards. Exp. Appl. Acarol. 67, 299–307. https://doi.org/10.1007/s10493-015-9950-z

Pung, O.J., Durden, L.A., Banks, C.W., Jones, D.N., 1994. Ectoparasites of Opossums and Raccoons in Southeastern Georgia. J. Med. Entomol.31, 915–919. https://doi.org/10.1093/jmedent/31.6.915

R Development Core Team, 2020. R: A Language and Environment for Statistical Computing.

Rainey, T., Occi, J.L., Robbins, R.G., Egizi, A., 2018. Discovery of *Haemaphysalis longicornis* (Ixodida: Ixodidae) parasitizing a sheep in New Jersey, United States. J. Med. Entomol. 55, 757–759. https://doi.org/10.1093/jme/tjy006

Raney, W.R., Perry, J.B., Hermance, M.E., 2022. Transovarial transmission of Heartland virus by invasive Asian longhorned ticks under laboratory conditions. Emerg. Infect. Dis. 28, 726–729. https://doi.org/10.3201/eid2803.210973

Reese, S.M., Petersen, J.M., Sheldon, S.W., Dolan, M.C., Dietrich, G., Piesman, J., Eisen, R.J., 2011. Transmission efficiency of *Francisella tularensis* by adult american dog ticks (Acari: Ixodidae). J. Med. Entomol. 48, 884–890. https://doi.org/10.1603/ME11005

Rochlin, I., Egizi, A., Ginsberg, H.S., 2022. Modeling of historical and current distributions of lone star tick, *Amblyomma americanum* (Acari: Ixodidae), is consistent with ancestral range recovery. Exp. Appl. Acarol. https://doi.org/10.1007/s10493-022-00765-0

Rochlin, I., Egizi, A., Narvaez, Z., Bonilla, D.L., Gallagher, M., Williams, G.M., Rainey, T., Price, D.C., Fonseca, D.M., 2023. Microhabitat modeling of the invasive Asian longhorned tick (*Haemaphysalis longicornis*) in New Jersey, USA. Ticks Tick Borne Dis. 14, 102126. https://doi.org/10.1016/j.ttbdis.2023.102126

Rosenberg, R., Lindsey, N.P., Fischer, M., Gregory, C.J., Hinckley, A.F., Mead, P.S., Paz-Bailey, G., Waterman, S.H., Drexler, N.A., Kersh, G.J., Hooks, H., Partridge, S.K., Visser, S.N., Beard, C.B., Petersen, L.R., 2018. *Vital Signs*⍰: Trends in Reported Vectorborne Disease Cases — United States and Territories, 2004–2016. MMWR Morb. Mortal. Wkly. Rep. 67, 496–501. https://doi.org/10.15585/mmwr.mm6717e1

Russart, N.M., Dougherty, M.W., Vaughan, J.A., 2014. Survey of ticks (Acari: Ixodidae) and tick-borne pathogens in North Dakota. J. Med. Entomol. 51, 1087–1090. https://doi.org/10.1603/me14053

Schmidt, K.A., Ostfeld, R.S., Schauber, E.M., 1999. Infestation of *Peromyscus* leucopus and *Tamias striatus* by *Ixodes scapularis* (Acari: Ixodidae) in relation to the abundance of hosts and parasites. J. Med. Entomol. 36, 749–757. https://doi.org/10.1093/jmedent/36.6.749

Schulze, T.L., Lakat, M.F., Bowen, G.S., Parkin, W.E., Shisler, J.K., 1984. *Ixodes dammini* (Acari: Ixodidae) and other ixodid ticks collected from white-tailed deer in New Jersey, USA. I. Geographical distribution and its relation to selected environmental and physical factors. J. Med. Entomol. 21, 741–749. https://doi.org/10.1093/jmedent/21.6.741

Shannon, C.E., 1948. A Mathematical Theory of Communication. The Bell System Technical Journal. 27, 379– 423.

Sharma, R., Cozens, D.W., Armstrong, P.M., Brackney, D.E., 2021. Vector competence of human-biting ticks *Ixodes scapularis, Amblyomma americanum* and *Dermacentor variabilis* for Powassan virus. Parasit. Vectors. 14, 466. https://doi.org/10.1186/s13071-021-04974-1

Shaw, M.T., Keesing, F., Mcgrail, R., Ostfeld, R.S., 2003. Factors influencing the distribution of larval blacklegged ticks on rodent hosts. Am. J. Trop. Med. Hyg. 68, 447–452. https://doi.org/10.4269/ajtmh.2003.68.447

Shaw, S.E., Day, M.J., Birtles, R.J., Breitschwerdt, E.B., 2001. Tick-borne infectious diseases of dogs. Trends Parasitol. 17, 74–80. https://doi.org/10.1016/S1471-4922(00)01856-0

Sidge, J.L., Foster, E.S., Buttke, D.E., Hojgaard, A., Graham, C.B., Tsao, J.I., 2021. Lake Michigan insights from island studies: the roles of chipmunks and coyotes in maintaining *Ixodes scapularis* and *Borrelia burgdorferi* in the absence of white-tailed deer. Ticks Tick Borne Dis. 12, 101761. https://doi.org/10.1016/j.ttbdis.2021.101761

Sivakumar, T., Tattiyapong, M., Okubo, K., Suganuma, K., Hayashida, K., Igarashi, I., Zakimi, S., Matsumoto, K., Inokuma, H., Yokoyama, N., 2014. PCR detection of *Babesia ovata* from questing ticks in Japan. Ticks Tick Borne Dis. 5, 305–310. https://doi.org/10.1016/j.ttbdis.2013.12.006

Smart, D.L., Caccamise, D.F., 1988. Population dynamics of the American dog tick (Acari: Ixodidae) in relation to small mammal Hosts. J. Med. Entomol. 25, 515–522. https://doi.org/10.1093/jmedent/25.6.515

Sonenshine, E. by D.E., Roe, R.M. (Eds.), 2013. Biology of Ticks Volume 1, Second Edition. ed. Oxford University Press, Oxford, New York.

Stanley, H.M., Ford, S.L., Snellgrove, A.N., Hartzer, K., Smith, E.B., Krapiunaya, I., Levin, M.L., 2020. The ability of the invasive Asian longhorned tick *Haemaphysalis longicornis* (Acari: Ixodidae) to acquire and transmit *Rickettsia rickettsii* (Rickettsiales: Rickettsiaceae), the agent of Rocky Mountain Spotted Fever, under laboratory conditions. J. Med. Entomol. 57, 1635–1639. https://doi.org/10.1093/jme/tjaa076

Telford III, S.R., Mather, T.N., Adler, G.H., Spielman, A., 1990. Short-tailed shrews as reservoirs of the agents of Lyme disease and human babesiosis. J. Parasitol. 76, 681. https://doi.org/10.2307/3282982

Thompson, A.T., Dominguez, K., Cleveland, C.A., Dergousoff, S.J., Doi, K., Falco, R.C., Greay, T., Irwin, P., Lindsay, L.R., Liu, J., Mather, T.N., Oskam, C.L., Rodriguez-Vivas, R.I., Ruder, M.G., Shaw, D., Vigil, S.L., White, S., Yabsley, M.J., 2020. Molecular characterization of *Haemaphysalis* species and a molecular genetic key for the identification of *Haemaphysalis* of North America. Front. Vet. Sci. 7, 141. https://doi.org/10.3389/fvets.2020.00141

Thompson, A.T., White, S.A., Shaw, D., Garrett, K.B., Wyckoff, S.T., Doub, E.E., Ruder, M.G., Yabsley, M.J., 2021. A multi-seasonal study investigating the phenology, host and habitat associations, and pathogens of *Haemaphysalis longicornis* in Virginia, U.S.A. Ticks Tick Borne Dis. 12, 101773. https://doi.org/10.1016/j.ttbdis.2021.101773

Tokarz, R., Sameroff, S., Tagliafierro, T., Jain, K., Williams, S.H., Cucura, D.M., Rochlin, I., Monzon, J., Carpi, G., Tufts, D., Diuk-Wasser, M., Brinkerhoff, J., Lipkin, W.I., 2018. Identification of novel viruses in *Amblyomma americanum*, *Dermacentor variabilis*, and *Ixodes scapularis ticks*. mSphere 3, e00614–17. https://doi.org/10.1128/mSphere.00614-17

Tryon, C.A., Snyder, D.P., 1973. Biology of the eastern chipmunk, *Tamias striatus*: life tables, age distributions, and trends in population numbers. J. Mammal. 54, 145–168. https://doi.org/10.2307/1378877

Tufts, D.M., Goodman, L.B., Benedict, M.C., Davis, A.D., VanAcker, M.C., Diuk-Wasser, M., 2021. Association of the invasive *Haemaphysalis longicornis* tick with vertebrate hosts, other native tick vectors, and tick-borne pathogens in New York City, USA. Int. J. Parasitol. 51, 149–157. https://doi.org/10.1016/j.ijpara.2020.08.008

Tufts, D.M., VanAcker, M.C., Fernandez, M.P., DeNicola, A., Egizi, A., Diuk-Wasser, M.A., 2019. Distribution, host-seeking phenology, and host and habitat associations of *Haemaphysalis longicornis* ticks, Staten Island, New York, USA. Emerg. Infect. Dis. 25, 792–796. https://doi.org/10.3201/eid2504.181541

USDA, U.S.D. of A., 2023. National Haemaphysalis longicornis (Asian longhorned tick) situation Report. United States Department of Agriculture, Beltsville, MD.

Watts, J., Playford, M., Hickey, K., 2016. *Theileria orientalis*: a review. New Zeal. Vet. J. 64, 3–9. https://doi.org/10.1080/00480169.2015.1064792

White, S.A., Bevins, S.N., Ruder, M.G., Shaw, D., Vigil, S.L., Randall, A., Deliberto, T.J., Dominguez, K., Thompson, A.T., Mertins, J.W., Alfred, J.T., Yabsley, M.J., 2021. Surveys for ticks on wildlife hosts and in the environment at Asian longhorned tick (Haemaphysalis longicornis)-positive sites in Virginia and New Jersey, 2018. Transbound. Emerg. Dis. 68, 605–614. https://doi.org/10.1111/tbed.13722

Wormser, G.P., McKenna, D., Piedmonte, N., Vinci, V., Egizi, A.M., Backenson, B., Falco, R.C., 2020. First recognized human bite in the United States by the Asian longhorned tick, *Haemaphysalis longicornis*. Clinical Infectious Diseases 70, 314–316. https://doi.org/10.1093/cid/ciz449

Yahner, R.H., 1978. The adaptive nature of the social system and behavior in the eastern chipmunk, *Tamias striatus*. Behav. Ecol. Sociobiol. 3, 397–427. https://doi.org/10.1007/BF00303202

Zanaga, D., Van De Kerchove, R., De Keersmaecker, W., Souverijns, N., Brockmann, C., Quast, R., Wevers, J., Grosu, A., Paccini, A., Vergnaud, S., Cartus, O., Santoro, M., Fritz, S., Georgieva, I., Lesiv, M., Carter, S., Herold, M., Li, Linlin, Tsendbazar, N.E., Ramoino, F., Arino, O., 2021. ESA WorldCover 10 m 2020 v100.

Zar, J.H., 2010. Two-sample hypotheses. testing for difference between two diversity indices, in: Biostatistical Analysis, Chapter. Pearson Prentice Hall, New Jersey, pp. 174–176.

Zhao, L., Li, J., Cui, X., Jia, N., Wei, J., Xia, L., Wang, H., Zhou, Y., Wang, Qian, Liu, X., Yin, C., Pan, Y., Wen, H., Wang, Qing, Xue, F., Sun, Y., Jiang, J., Li, S., Cao, W., 2020. Distribution of *Haemaphysalis longicornis* and associated pathogens: analysis of pooled data from a China field survey and global published data. Lancet Planet. Health 4, e320–e329. https://doi.org/10.1016/S2542-5196(20)30145-5

Zheng, H., Yu, Z., Zhou, L., Yang, X., Liu, J., 2012. Seasonal abundance and activity of the hard tick *Haemaphysalis longicornis* (Acari: Ixodidae) in North China. Exp. Appl. Acarol. 56, 133–141. https://doi.org/10.1007/s10493-011-9505-x

Zimmerman, R.H., McWherter, G.R., Bloemer, S.R., 1988. Medium-sized mammal hosts of *Amblyomma americanum* and *Dermacentor variabilis* (Acari: Ixodidae) at land between the lakes, Tennessee, and effects of integrated tick management on host infestations. J. Med. Entomol. 25, 461–466. https://doi.org/10.1093/jmedent/25.6.461

