## Supplementary figures and images for "Ticks (Acari: Ixodida) on synanthropic small and medium-sized mammals in areas of the northeastern United States infested with the Asian longhorned tick, *Haemaphysalis longicornis*"

### Figure S1

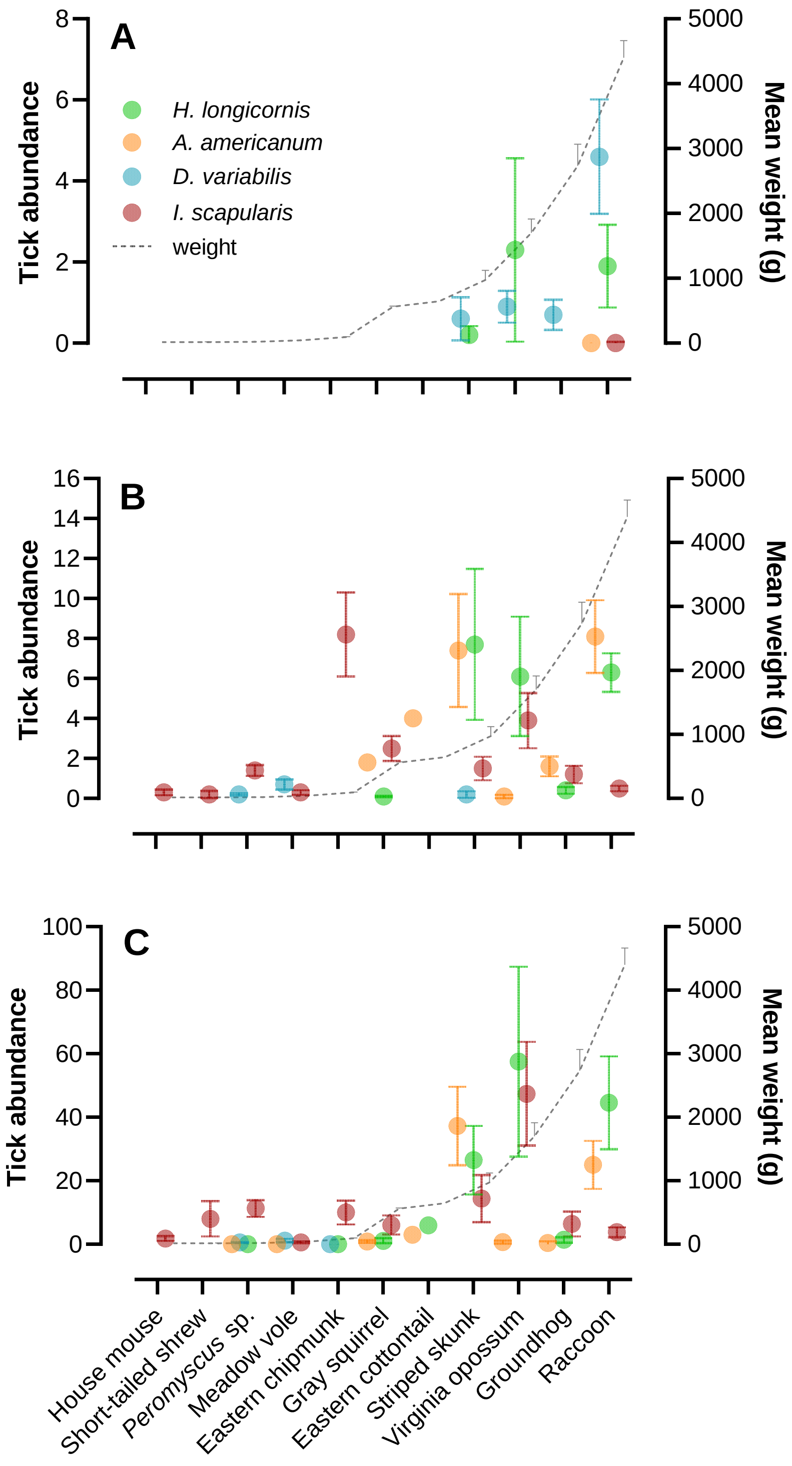
